# Protein Phosphorylation Orchestrates Acclimations of *Arabidopsis* Plants to Environmental pH

**DOI:** 10.1101/2023.03.19.533375

**Authors:** Dharmesh Jain, Wolfgang Schmidt

**Affiliations:** Molecular and Biological Agricultural Sciences Program, Taiwan International Graduate Program, Academia Sinica and National Chung-Hsing University, Taipei 11529, Taiwan; Graduate Institute of Biotechnology, National Chung-Hsing University, Taichung 40227, Taiwan; Institute of Plant and Microbial Biology, Academia Sinica, Taipei 11529, Taiwan; Biotechnology Center, National Chung-Hsing University, Taichung 40227, Taiwan; Genome and Systems Biology Degree Program, College of Life Science, National Taiwan University, Taipei 10617, Taiwan

## Abstract

Environment pH (pH_*e*_) is a key parameter that dictates a surfeit of conditions critical to plant survival and fitness. To elucidate the mechanisms that recalibrate cytoplasmic and apoplastic pH homeostasis, we conducted a comprehensive proteomic/phosphoproteomic inventory of plants subjected to transient exposure to acidic or alkaline pH, an approach that covered the majority of protein-coding genes of the model plant *Arabidopsis thaliana*. Our survey revealed a large set so far undocumented pH_*e*_-dependent and potentially pH-specific phospho-sites, indicative of extensive post-translational regulation of proteins involved in the acclimation to pH_*e*_. Changes in pH_*e*_ altered both electrogenic H^+^ pumping via P-type ATPases and H^+^/anion co-transport processes, leading to massively altered net trans-plasma membrane translocation of H^+^ ions. In pH 7.5 plants, transport (but not the assimilation) of nitrogen via NRT2-type nitrate and AMT1- type ammonium transporters was induced, conceivably to increase the cytosolic H^+^ concentration. Exposure to acidic pH resulted in a marked repression of primary root elongation. No such cessation was observed in *nrt2.1* mutants, suggesting a role of NRT2.1 in negatively regulating root growth in response to acidic pH. Sequestration of iron into the vacuole via phosphorylation and abundance of the vacuolar iron transporter VTL5 was inversely regulated in response to high and low pH_*e*_, presumptively in anticipation of changes in iron availability associated with alterations of pH_*e*_ in the soil. A pH-dependent ‘phospho-switch’ was also observed for the ABC transporter PDR7, suggesting changes in activity and, possibly, substrate specificity. Unexpectedly, the effect of pH_*e*_ was not restricted to roots and provoked pronounced changes in the leaf proteome. In both roots and shoots, the plant-specific TPLATE complex components AtEH1 and AtEH2 – essential for clathrin-mediated endocytosis – were differentially phosphorylated at multiple sites in response to pH_*e*_, indicating that the endocytic cargo protein trafficking is orchestrated by pH_*e*_.

## INTRODUCTION

In lieu of strategies to escape unfavorable conditions, plants have evolved mechanisms to cope with and adapt to the specific circumstances of their natural habitat. A key role in defining whether or not a given species is able to thrive in a given soil is played by the proton concentration – or pH – of the soil solution. Soil pH dictates a plethora of processes that are crucial for plant fitness, including but not limited to the availability of essential mineral nutrients, the level of potentially toxic ions in the soil solution, and the composition and activity of the soil microbiome. As a rule, acid soils (i.e., soils featuring pH values below 5.5) are associated with elevated Fe^2+^ and Mn^2+^ concentrations, decreased nitrification (and subsequent low levels of nitrate), and leaching of cations such as Ca^2+^, Mg^2+^, and K^+^ (1). The biggest constraint posed on plants thriving in acid soils is the high concentration of exchangeable Al ions, in particular the potentially toxic Al^3+^ species (2). Alkaline soils, on the other hand, often contain abundant levels of bicarbonate (the referred to as calcareous soils), which efficiently buffers soil pH and counteracts attempts to increase nutrient availability by root-mediated acidification of the rhizosphere. Species thriving in such soils are referred to as calcicole (Greek for chalk-loving), as opposed to calcifuge (chalk-fleeing) species. Typically, alkalinity restricts the availability of several essential mineral nutrients, foremost iron, manganese, and phosphorus (3).

Unfavorable soil pH compromises both the physiological performance and fitness of the plant, but the reasons for growth restrictions may differ between acid and alkaline soils.

Generally, low pH favors growth by relaxation of wall stress (acid growth theory; (4)), supporting cell elongation by the induction of cell-wall loosening enzymes (5). However, unrestricted growth is not necessarily advantageous in acid soils. High levels of ammonium or Al^3+^ ions lead to reduced root elongation through accumulation of auxin in the root transition zone (6), a response that protect the plant from excessive accumulation of potentially toxic ions. By contrast, alkaline pH restricts loosening of the cell wall and, thus, compromises plant growth. For plants that are not well adapted to such conditions, slow growth might be advantageous to avoid depletion of essential mineral nutrients that have limited mobility in alkaline soils.

Using Arabidopsis as a model, Liu et al. (7) recently showed that plants perceive and process information on the pH of the root apoplast (pH_*apo*_) via a bimodal peptide-receptor system. At low pH, protonation of a sulfotyrosine residue of the Root Growth Factor (RGF) peptide allows formation of a complex comprising RGF, its receptor RGFG, and the coreceptor SERK. This complex promotes meristem development by tuning gradients of the transcription factor PLETHORA, a key regulator of root development (8, 9). At alkaline pH, RGF is deprotonated, which destabilizes the RGF-RGFR-SERK complex. Instead, high pH_*apo*_ is associated with PAMP-triggered immunity, induced by a separate component of the pH-sensing system, comprising the peptide Pep, its receptor PEPR, and the coreceptor BAK1 (7). Thus, pH serves as a critical checkpoint to balance two crucial tasks of roots, growth and defense.

How plants recalibrate cytosolic and apoplastic proton homeostasis in response to alterations in external pH (pH_*e*_) is not well explored. Two mechanisms, the biophysical pH stat, i.e., the electrogenic transport of protons across the plasma membrane, and the biochemical pH stat, i.e., changes in cytosolic proton concentration via protonation and deprotonation of organic acids, have been proposed as main mechanisms controlling cytosolic pH (pH_*cyt*_) (10), but these concepts might not be sufficient to cover the complexity of intracellular and extracellular pH control. Exposure to both high and low pH_*e*_ was shown to cause pronounced changes in root transcriptional profiles (11, 12). Transcriptional changes are, however, not always an adequate blueprint for changes in protein abundance and, even less so, for the physiological readout of gene activity (13), suggesting that cataloging the abundance of proteins is more likely to reflect relevant gene activity than charting transcriptomic profiles. The caveat of bottom-up (‘shotgun’) proteomics approaches is the generally insufficient coverage of the proteome, owing to the difficulty of capturing lowly abundant or instable proteins. This impediment has been – to a large extent – ironed out by advanced mass spectrometry technologies and improved gene annotations, allowing for comprehensive proteomic surveys that are not restricted to a particular population of cellular proteins. Here, we provide an exhaustive proteomic inventory of roots and shoots of the model plant *Arabidopsis thaliana* subjected to short-term exposure to high or low media pH, an approach that covered more than 60% of the protein-coding genes. To unveil signal transduction cascades that involve protein phosphorylation, we introduced a phosphoprotein enrichment step that captured a large number of novel, putatively pH-specific phosphorylation sites. Our analysis revealed an elaborate interplay between processes dictating growth, defense, and the control of cytoplasmic pH, responses that are orchestrated by regulating gene activity at different levels.

## EXPERIMENTAL PRECEDURES

### Plant Growth

Seeds of Arabidopsis (*Arabidopsis thaliana* (L.) Heynh, accession Col-0) were purchased from the Arabidopsis Biological Resource Center (ABRC; Ohio State University). Seeds of the *nrt2.1* mutant (Salk_141712) were provided by Dr. Yi-Fang Tsay (IMB, Academia Sinica, Taipei Taiwan). Plants were grown under sterile conditions in a growth chamber on agar-based media as described by Estelle and Somerville (14) with slight modifications. Seeds were surface-sterilized by immersion in 30% (v/v) commercial bleach containing 6% NaClO and 70% (v/v) absolute ethanol containing 0.1% (v/v) TWEEN 20 for 6 min, followed by five rinses with absolute ethanol. The growth medium was comprised of 5 mM KNO3, 2 mM MgSO4, 2 mM Ca(NO3)2, 2.5 mM KH2PO4, 40 µM Fe^3+^-EDDHA, 70 µM H3BO3, 14 µM MnCl2, 1 µM ZnSO4, 0.5 µM CuSO4, 0.01 µM CoCl2, and 0.2 µM Na2MoO4, supplemented with 1.5% (w/v) sucrose and solidified with 0.5% Gelrite pure (Kelco). For control and acidic pH, 1 g/L MES was added, and the pH was adjusted to 4.5 (acidic) and 5.5 (control) with KOH. For alkaline media, 1 g/L MOPS was added, and the pH was adjusted to 7.5 with KOH. For growth analysis, seeds were directly sown on media adjusted to different pH values and stratified for 3 days at 4 °C in the dark before the plates were transferred to a growth chamber and grown at 22 °C under continuous illumination (50 µmol m^−2^ s^−2^) for 14 days and collected for growth parameter analysis. For proteomics and phosphoproteomics analysis, seeds were directly sown on pH 5.5 media and stratified for 3 days at 4 °C in the dark before the plates were transferred to a growth chamber and grown at 22 °C under continuous illumination (50 µmol m^−2^ s^−2^). After 14 days, seedlings were carefully transferred to media adjusted to either pH 4.5, 5.5, or 7.5 for 6 hours. Then, samples were collected and transferred to liquid nitrogen at at the end of the experimental period.

### Root Length, Rosette Size, and Dry Weight

For morphological analysis, plants were grown (vertically for root length, and horizontally for rosette diameter) on media with pH adjusted to either 4.5, 5.5, or 7.5 for 14 days. Seedlings were imaged with a digital camera (Alpha 7 IV, Sony). Primary root length and rosette size was analyzed using the ImageJ software. After imaging, root and shoot samples were collected (20 specimen per replicate) and dried in an oven at 45 °C for two days. Results were expressed as means ± standard error (SE). All measurements indicate the average of three independent biological experiments.

### Chlorophyll Concentration

Seedlings were grown for 14 days continuously on media adjusted to various pH values. Five shoot samples were collected, weighed, and immediately placed in an Eppendorf tube with steel beads, immersed in liquid nitrogen and stored in −80 °C until analyzed. For analyzing chlorophyll concentration, samples were homogenized using a TissueLyzer II (Qiagen).

Chlorophyll was extracted and measured following a protocol from Mackinney (15). Frozen tissues were dissolved in 500 μL 80% (v/v) acetone. The tubes were centrifuged at 13,200 rpm for 5 minutes and the supernatant was collected in the new amber Eppendorf tube. The 80% acetone step was repeated thrice or until the pellet became white and all the extracts were pooled together. Finally, the pooled extract was mixed thoroughly and centrifuged again to pellet down the remaining cell debris. Two hundred μL of extract was used for measuring the absorbance at 663, 647, and 750 nm in a PowerWave XS2 microplate spectrophotometer from BioTek.

Absorbance at 750 nm was used to correct absorbance at 663 and 647 nm. The equation described by Lichtenthaler (16) was used for calculating the chlorophyll concentration.

### Soil Experiments

Soil with a pH of 5.6 was used for control experiments. For preparing alkaline substrate, 20 g kg^−1^ CaCO3 and 12 g kg^−1^ NaHCO3 were added to the soil and mixed thoroughly, leading to a pH of 7.2. Col-0 seeds were stratified for 3 days at 4° C in the dark before they were sown on the soil. Pots were transferred to a growth chamber and plants were grown at 22 °C/18 °C and 16 h/8 h light/dark regime at a light intensity of 120 µmol m^−2^ s^−1^. To maintain iron available for plants, 2.0 mL of a 4.4 g L^−1^ Fe-sequestrene (6% Iron Chelate, PlantMedia) solution was added to each pot (17). Twenty-one days after sowing, pictures of the rosette phenotype were taken. Rosette size was quantified using the ImageJ software. All measurements report the average of four independent biological experiments, each involving five seedlings. For the determination of silique length and seed number, plants were grown until maturation and first four developed siliques were collected from the primary inflorescence. Collected siliques were than photographed using Dissecting fluorescent microscope (Lumar V12, Zeiss, Germany). Silique length and seed number per silique were analyzed using ImageJ software. For seed weight, plants were grown until maturation. Seed weight was determined in four independent biological replicates.

### Flower Morphology

To determine pollen viability, pollen number, and anther size, mature unopened flower buds with indehiscent anthers (anther stage 12-13) were collected and stained in Alexander stain overnight at 4 °C (18). Stained flower buds were collected and carefully placed on a microscope slide. Sepal and petals were removed with the help of a thin needle. Style and filament morphology was photographed using a dissecting fluorescent microscope (Lumar V12, Zeiss, Germany). Anthers were collected by cutting the filament and putting them on a separate slide with a coverslip. The coverslip was gently pressed until individual pollen grains were visible.

Anthers were then photographed using a compound microscope (Z1 Imager, Zeiss, Germany). Pollen number was quantified using the ImageJ software (National Institutes of Health). Pollen number was determined in four independent biological replicates.

### Proteomics and Phosphoproteomics Analysis

Proteomics and phosphoproteomics analyses were performed as described previously (19) with slight modifications. For analysis, 1 g of root or shoot tissues was coarsely homogenized with piston and mortar and carefully transferred to a 15 mL Falcon tube. One mL freshly prepared 5% SDS containing 2x Roche cOmpleteTM, EDTA-free protease inhibitor cocktail, 1X phosphatase inhibitor cocktail 3, and 1X phosphatase inhibitor cocktail 2 was added to the tubes, vortexed, sonicated on ice with 15 s on /15s off cycles and then boiled at 95 °C for 5 min. Samples were then placed on ice for 10 min and centrifuged at 5,000 g at 4 °C for 60 min.

The supernatants (plant lysates) were collected and transferred to new low protein-binding collection tubes. Subsequently, 150 μL plant lysate solutions were transferred into a fresh 1.7 mL tube and 600 μL 100% methanol and 150 μL 100% chloroform were added. After each of these steps, the samples were vortexed and spun down. Then, 450 μL ddH2O were added to the tubes, the samples were vortexed and centrifuged at 16,000 g for 3 mins. After centrifugation, the upper aqueous layer was discarded and 600 μL 100% methanol were added into the tubes. The content was mixed carefully with the help of a pipette and then centrifuged at 16,000 g for 3 mins. This step was repeated once. Finally, the supernatant was discarded and the protein pellets were air-dried. The dried protein pellets were resuspended in 6 M urea and 50 mM Tris-HCl, pH 8.5.

Protein concentrations were quantified with a PierceTM 660 nm protein assay following the manufacturer’s protocol. The protein was then diluted to 4 μg/μL using 6 M urea. Next, 100 mM TCEP (Tris 2-carboxyethyl phosphine hydrochloride; Sigma-Aldrich, C4706) were added to a final concentration of 10 mM followed by adding CAA (800 mM 2-chloroacetamide; Sigma-Aldrich, C0267) to a final concentration of 40 mM.

For in-solution trypsin digestion, the protein concentrates (in 6 M urea) were diluted 4- fold (to 1.5 M urea) using 50 mM TEAB (triethylammonium bicarbonate; Sigma-Aldrich; Cat. No. T7408). The protein was digested using lysyl endopeptidase (Lys-C) in a 1:50 enzyme-to-substrate (E/S) ratio (2 μg enzyme and 100 μg protein) for 4 h at RT (room temperature).

Subsequently, modified trypsin at a 1:50 E/S ratio was added and incubated at 37 °C overnight. Then, 10% TFA was added to a final concentration of 1% to acidify the solution. A solution pH of 2-3 was determined using pH indicator paper. Next, the samples were desalted on a C18 solid-phase extraction cartridge. The cartridge were washed 4 times using 0.1% TFA, followed by centrifugation at 100 g for 1 min at RT. Finally, the peptides were eluted by 0.1% TFA in 75% ACN. Eluates were dried with a SpeedVac vacuum concentrator and stored at −80 °C until further analysis.

In order to pursue tandem mass tag (TMT) labelling, 100 μg digested peptide samples (dried pellets) were resuspended in 100 μL HEPES buffer (200 mM, pH 8.5). TMT reagents were reconstituted in 40 μL anhydrous acetonitrile, and peptides were labelled with the TMT reagents (Thermo Fisher Scientific) according to the manufacturer’s protocol for 1 h at RT. Then, samples were quenched by adding 5% hydroxylamine, incubated for 15 minutes, mixed in a 5 mL tube, and acidified to a final concentration of 1% TFA. The solution was later desalted using a C18 cartridge and centrifuged at 100 g for 1 min at RT. The cartridge was washed 4 times with 0.1% TFA and centrifuged at 100 g for 1 min at RT. Peptides were eluted using 75% ACN and dried using a SpeedVac.

*Tip Preparation for Phospho-Enrichment -* A polypropylene frit was inserted into a 200 μL pipette tip. Ten mg Ni-NTA beads were suspended with 400 μL 6% acetic acid (AA), pH 3.0. The bead solution was loaded into the IMAC StageTip and the StageTip was centrifuged at 200 g for 3 mins to remove the solution. Then, 100 μL EDTA (50 mM) was loaded onto the IMAC StageTip and centrifuged at 200 g for 3 min. Next, 100 μL AA (6%) was loaded onto the IMAC StageTip and centrifuged at 200 g for 3 min. Then, 100 μL FeCl3 (50 mM) in 6% AA was loaded onto the IMAC StageTip and centrifuged at 200 g for 3 min. Next, 100 μL 6% AA (pH 3.0) were loaded onto the IMAC StageTip and centrifuged at 200 g for 3 min. The dried and labeled peptides were suspended in 100 μL 6% AA (pH 3.0), loaded onto the IMAC StageTip and centrifuged at 200 g for 3 min. After this step, the flowthrough was processed separately for total proteomic analysis, and the filtrate was processed separately for phosphoproteomics.

*IMAC StageTip Processing for Phosphophoproteomics -* The IMAC StageTip was washed with 100 μL 4.5% AA/25% ACN, centrifuged at 200 g for 3 min, washed with 100 μL of 6% AA and centrifuged at 200 g for 3 min. A C18 StageTip with a layer of polypropylene frit was prepped following the manufacturer’s instructions. One mg C18 beads was suspended in 100 μL methanol (100%). The StageTip was centrifuged at 1,000 g for 5 min in order to pass through the solution. Next, 20 μL 200 mM NH4HCO2 (20%) and 80% ACN (pH 10.0) were added to the C18 StageTip and centrifuged at 1,000 g for 2 min. Then, 20 μL NH4HCO2 (200 mM, pH 10.0) were added and the StageTip was centrifuged at 1,000 g for 5 min. The washed IMAC StageTip was placed inside the C18 StageTip. Next, 100 μL NH4H2PO4 (200 mM, pH 4.4) were added to the IMAC StageTip and the solution was allowed to pass through the two layers of the StageTip by centrifugation at 500 g for 10 min at RT. After that, the IMAC StageTip was discarded. Then 20 μL NH4HCO2 (200 mM) were added and the C18 StageTip was centrifuged at 1,000 g for 2 min at RT. Next 20 μL NH4CO2 (91% 200 mM) and 9% ACN (pH 10.0) were added to the elute the bound phosphopeptides as fraction 1 and centrifuged at 1,000 g for 2 min at RT. These steps were repeated five times to yield six separate phosphopeptide fractions, in each step the percentage of 200 mM NH4CO2 was decreased as follows: 88%, 85%, 82%, 79%, 20%, and the percentage of ACN (pH 10.0) was changed to 12%, 15%, 18%, 21%, 80% at the respective steps. The eluates were dried using a SpeedVac. The samples were stored at −80 °C.

*Tip Preparation for Total Proteome Analysis -* The C18 StageTip was prepared with a layer of polypropylene frit. One mg C18 beads was suspended in 100 μL 100% methanol. These suspended beads were added to the tip and the tip was centrifuged at 1,000 g for 2 min at RT. After that, 20 μL NH4HCO2 (20% 200 mM) and 80% ACN (pH 10.0) were added to the C18 StageTip and centrifuged at 1,000 g for 2 min at RT. Then, 20 μL NH4HCO2 (200 mM NH4HCO2, pH 10.0) were added and the C18 StageTip was centrifuge at 1,000 g for 2 min at RT.

*Flowthrough Processing for Total Proteome Analysis -* After collecting, the flowthrough (IMAC FT peptides) was dried using a SpeedVac and dissolved in 100 μL 0.1% TFA. The concentration was measured and 10 μg IMAC FT peptides were diluted with 100 μL 0.1 % TFA. The solution was then added to a C18 StageTip and centrifuged at 1,000 g for 2 min at RT. Then, 20 μL of a solution containing 86% 200 mM NH4HCO2 and 14% ACN (pH 10.0) were added to the C18 StageTip and centrifuged at 1,000 g for 2 min to elute the bound peptide as fraction 1. This step was repeated five more times to yield six separate phosphopeptide fractions, in each step the percentage of 200 mM NH4HCO2 was decreased as follows: 83%, 80%, 77%, 74%, 20% and and the percentage of ACN (pH 10.0) was changed to 17%, 20%, 23%, 26%, 80% at the respective steps. The eluates were further dried using a SpeedVac. Samples were stored at −80°C.

### Liquid Chromatography

Liquid chromatography was performed on an EASY-nLC 1200 system (Thermo Fisher Scientific) coupled with an Orbitrap Fusion Lumos Tribrid mass spectrometer (Thermo Fisher Scientific) equipped with a Nanospray Flex Ion Source. The peptides of each fraction were dissolved in solvent A (0.1% formic acid in water) and centrifuged at 20,000 g for 10 min. The supernatant (peptide mixtures) was loaded onto the LC-MS/MS. The samples were separated using a segmented gradient for 120 min from 5% to 40% solvent B (acetonitrile with 0.1% formic acid) at a flow rate of 300 nL/min. The samples were maintained at 8 °C in an autosampler. The mass spectrometer was operated in positive ionization mode.

### Protein Identification

In order to identify and quantify proteins, Proteomic Discoverer 2.5 (Thermo Fisher Scientific) including the Mascot and SEQUEST software packages was used for database search. Searches were made against *Arabidopsis* protein databases in Araport11, and concatenated with a decoy database containing the randomized sequences of the original database. For each technical replicate, spectra from all fractions were combined into one MGF (Mascot generic format) file after loading the raw data, and MGF files were used to query protein databases. For each biological repeat, spectra from the three technical replicates were combined into one file and searched. The search parameters were as follows: trypsin was selected as an enzyme with two missed cleavages allowed; modifications of carbamidomethylation at Cys; TMT at peptide N-terminus and Lys; variable modifications of oxidation at Met; and phosphorylation at Ser, Thr, and Tyr were fixed; peptide tolerance was set to 10 ppm and MS/MS tolerance was set to 0.02 Da. Peptide charge was set to *Mr*, and monoisotopic mass. TMT10plex was chosen for quantification during the search consecutively. The search results were passed through additional filters before exporting the data files. For protein identification, the filters were set as follows: significance threshold *p* < 0.05 (with 95% confidence) and ion score or expected cutoff > 0.05 (with 95% confidence). For protein quantitation, filters were set as follows: ‘weighted’ was selected for protein ratio type; minimum precursor charge was set to 1, and minimum peptides was set to 2; only unique peptides were used to quantify proteins. Summed intensities were set as normalization, and outliers were removed automatically. The peptide threshold was set as above for homology.

### Bioinformatic Analyses

For the detection of new phosphorylation site, full experimental data sets were downloaded from the PhosPhat data base (20, 21). Next, the accession ID, Pubmed ID, and phosphosites were extracted and matched with the phosphopeptide files acquired in this study. Data extraction and matching was done using Perl scripts. The figure depicting the percentage of documented phosphosites was generated using the GraphPad prism software (v. 9; GraphPad Software, San Diego, CA, USA). Heatmaps were generated using CHM Builder (22); pathway analysis was performed using KEGG pathway analysis tool (23). Intracellular localization of proteins was inferred from the SUBA5 online web-based tool (24, 25). Transmembrane domain structures of the proteins were designed by mining experimental data and the web-based transmembrane prediction tool DeepTMHMM (26).

## RESULTS

### pHe Affects Growth and Fitness of Arabidopsis Col-0 Plants

To investigate the influence of pH_*e*_ on the growth of *A. thaliana*, we grew plants on media with strongly acidic (4.5), mildly acidic (5.5), and slightly alkaline (7.5) pH for a period of two weeks. Growth was optimal at pH 5.5 (Fig. 1). Alkalinity strongly affected shoot growth, shoot and root dry weight, while root length was only slightly (but statistically significant) affected. By contrast, growth on pH 4.5 media did not change rosette size or shoot dry weight, but dramatically reduced root growth (Fig. 1).

**FIG. 1.**
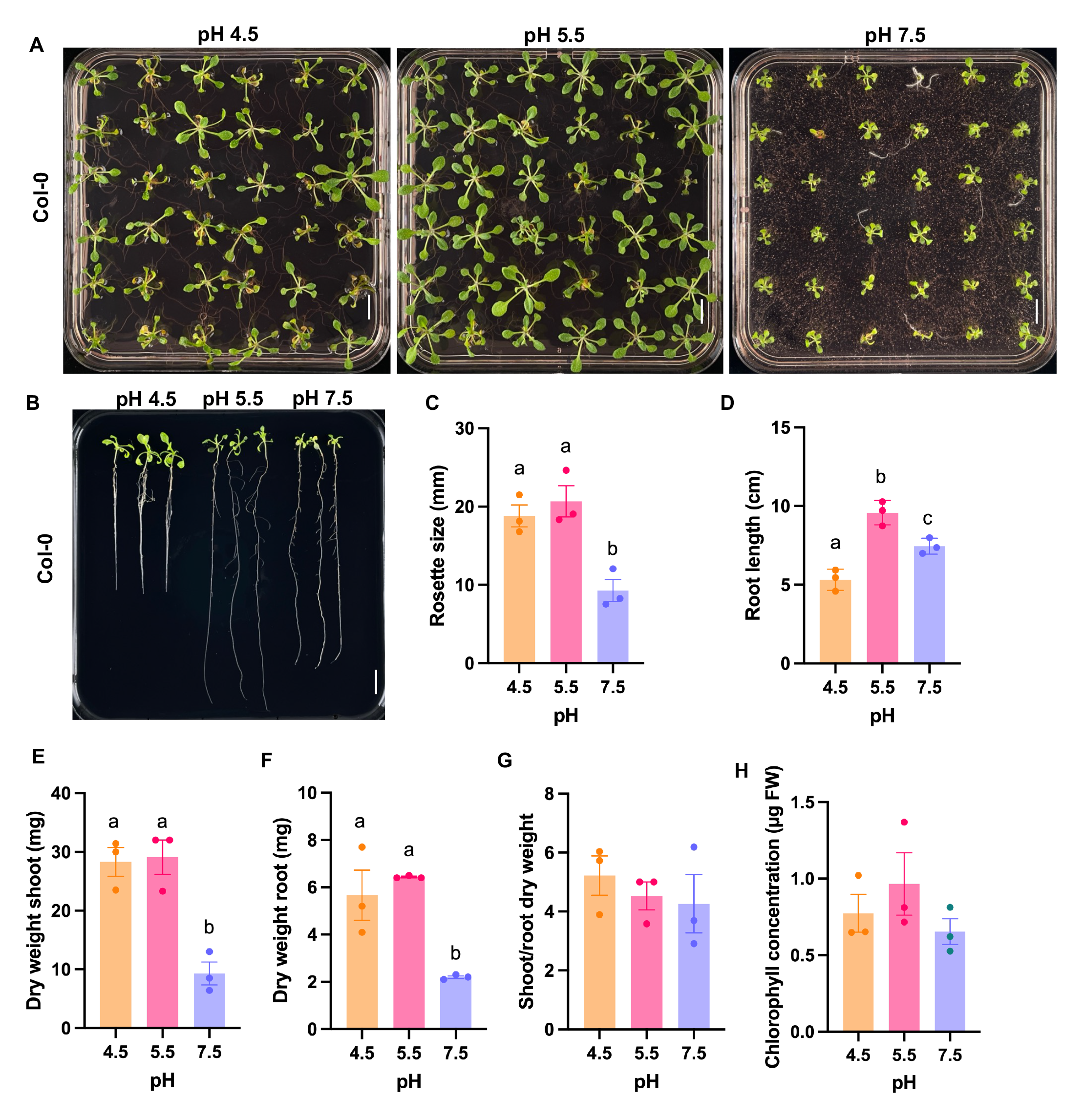
Phenotypes of *Arabidopsis* plants grown on media adjusted to different pH. *A*, leaf morphology. *B*, root phenotype. *C*, rosette diameter. *D*, root length. *E*, shoot dry weight. *F*, root dry weight. *G*, shoot-root ratio (dry weight). *H*, Chlorophyll concentration. 14-day-old seedling were used. Error bar represents the mean ± SE of four biologically independent experiments. Different letters indicate significant differences between averages using one-way ANOVA followed by *Tukey* post-hoc *test* (*p* < 0.05). Scale bar = 1cm.

Growing plants in slightly alkaline soil (pH 7.2), dramatically reduced plant size when compared with plants grown in soil with a pH that is optimal for *Arabidopsis* (pH 5.6) (Fig. 2*A*).

**FIG. 2.**
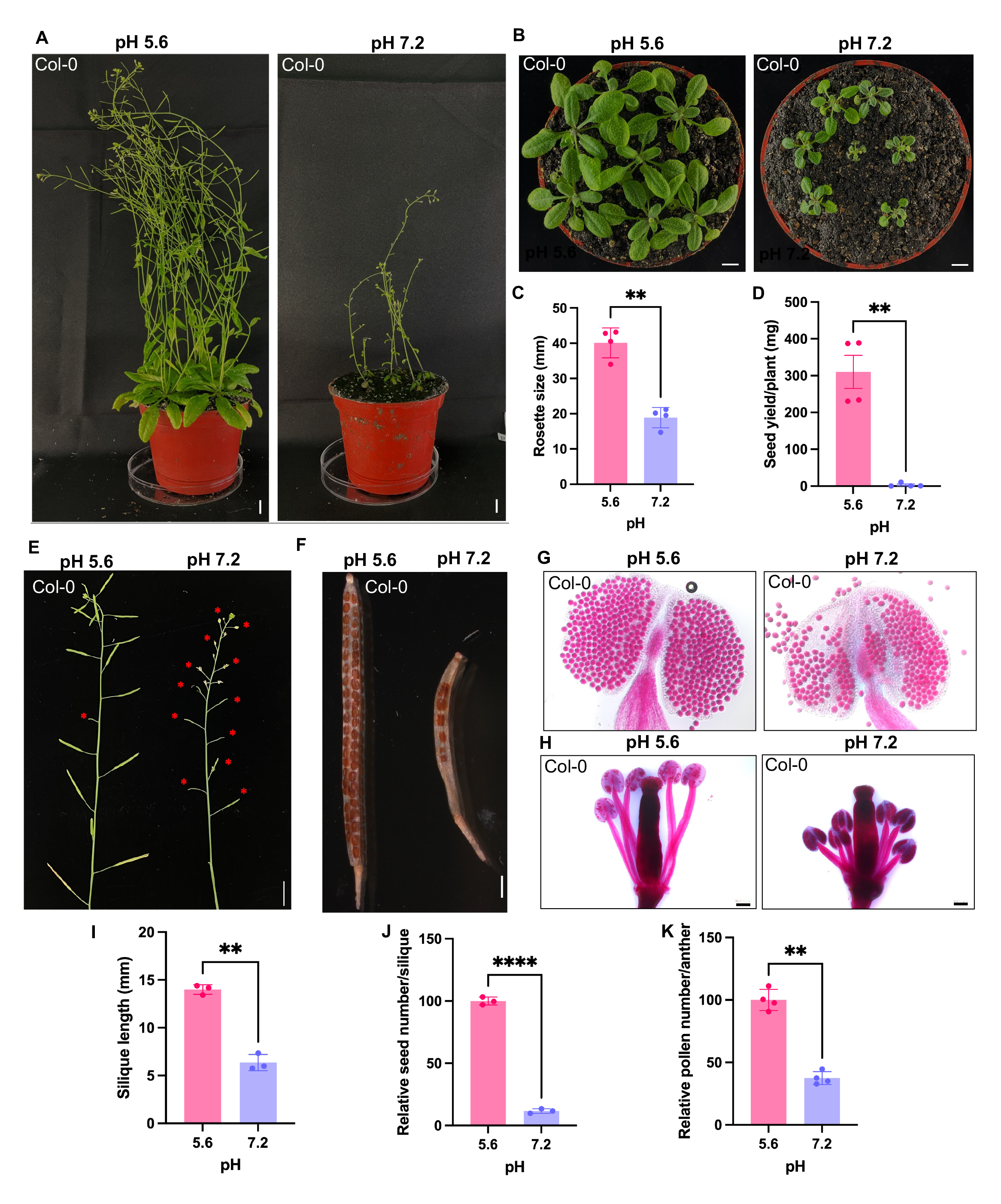
Morphological characterization of soil-grown *Arabidopsis* plants. *A*, phenotypes of 5- week-old plants grown in slightly acidic (left) and alkaline soil (right). *B*, rosette morphology of 21-day-old seedlings plants in slightly acidic (left) and alkaline soil (right). *C*, quantification of rosette size. *D*, seed yield. *E*, primary inflorescence of 5-week-old plants; red asterisk indicate aborted flower buds. *F*, mature silique of control plants (left) and of plants grown in pH 7.5 soil (right) showing aborted buds. *G*, Alexander’s staining showing the viability of pollens and anther size. *H*, morphology of filament and style. *I*, silique length. *J*, relative seed number per silique. K, quantification of the relative number of mature pollen grains. Error bar represents the mean ± SE of three or four biologically independent experiments. Statistical testing was carried out using Student’s t test. Asterisks indicate significant differences from the pH 5.6 control. **, *p* ≤ 0.01; ***, *p* ≤ 0.0001.

Similar to what has been observed for plants grown on plates, rosette size was reduced to approximately half of that of control plants grown at slightly acidic (pH 5.6) soil (Fig. 2*B*, *C*). In addition, seed yield was negligible at this pH, indicating severely reduced fitness of *Arabidopsis* Col-0 plants in alkaline soils (Fig. 2*D*). Growing plants on circumneutral soil altered the morphology of all reproductive parts of the plant, resulting in a dramatic reduction in seed yield (Fig. 2*D*). Primary inflorescences showed a high number of aborted flower buds (Fig. 2*E*).

Moreover, pollen number and filament size were reduced in plants grown in pH 7.2 soil, compromising self-pollination (Fig. 2*G*, *H*, *K*). Mature silique produced only a very small number of seeds under alkaline pH conditions (Fig. 2*F*, *I*, *J*). These data indicate that in addition to impaired growth, alkaline soil strongly affects the fitness of the Col-0 accession of *A. thaliana*.

### A Concatenated Approach to Map Proteins and Phosphorylation Events

To gain insights into processes governing the adaptation of *Arabidopsis* plants to alterations in pH_*e*_, we subjected plants precultured at optimal pH (5.5) to short-term (6 h) growth periods at either optimal, acidic, or alkaline pH. Subsequently, roots and shoots were subjected to an integrated proteomics/phosphoproteomics approach in which phosphopeptides and non-phosphopeptides were quantified from the same samples. The workflow of the experiment is shown in Figure 3. Proteins were extracted, precipitated, digested, and quality checked by Coomassie blue staining. After desalting, proteins were TMT-labeled and phosphopeptides were enriched by a spin-column-based strategy with agarose Ni-NTA beads. The flowthrough, containing TMT-labelled non-phosphopeptides, was subjected to high pH reversed-phase fractionation prior to LC-MS/MS analysis. Similarly, phosphopeptides were eluted from the column and sequenced alongside non-phosphopeptides. Detection and quantification of proteins was based on TMT ion reporter intensities recorded on an Thermo Q Exactive HF orbitrap mass spectrometer. Finally, MS/MS data from both fractions were analyzed using the Proteome Discoverer software.

**FIG. 3.**
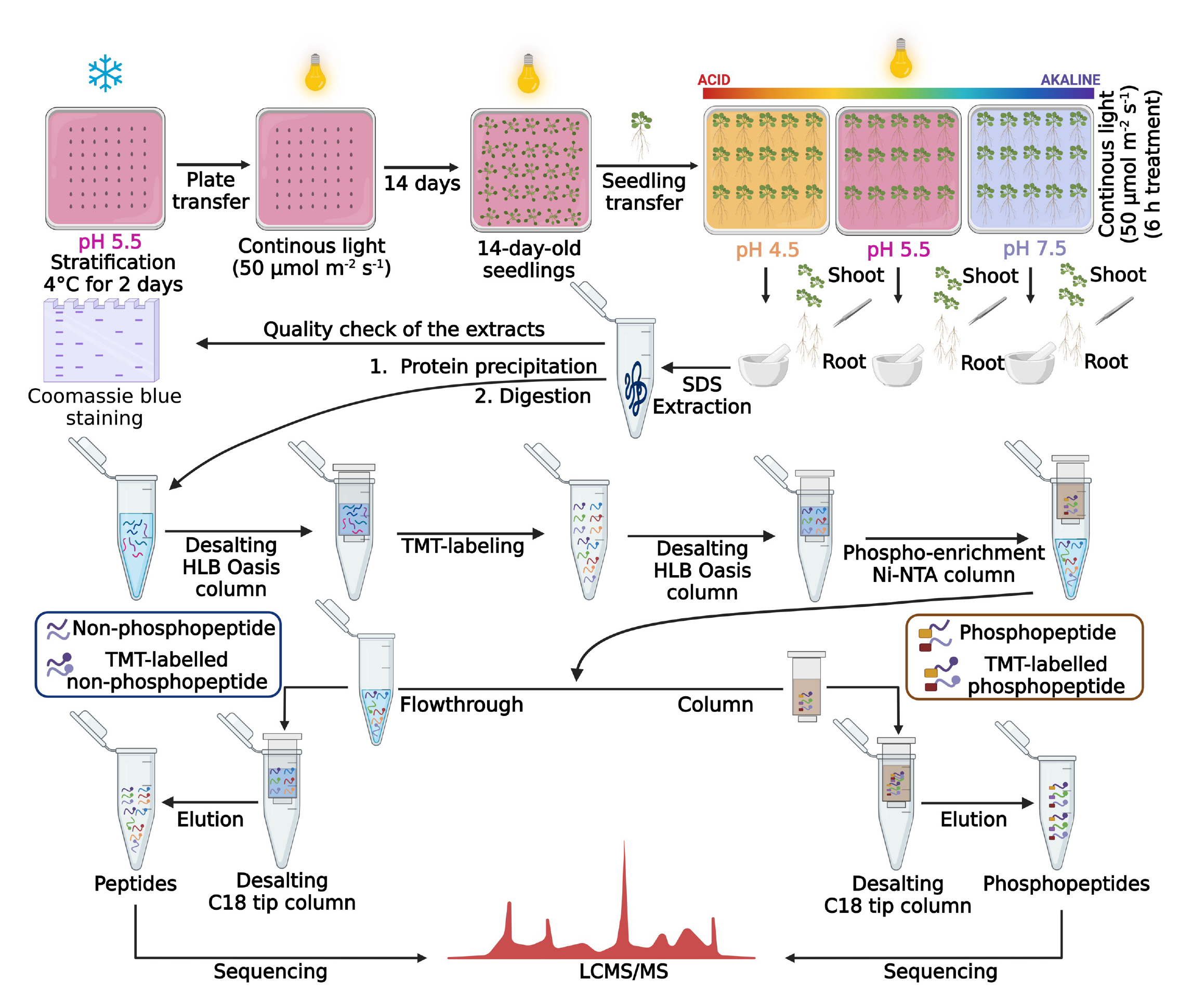
**Experimental workflow of the proteomics/phospho-proteomics analysis**. 14-day-old seedlings were grown for a 6 h period on media adjusted to pH 4.5, pH 5.5, and pH 7.5 and subjected to proteomics analysis. The experiments were performed in 3 biologically independent sets. Plants grown at pH 5.5 were used as a control. Details are given in the Materials and Methods section; a detailed protocol has been published in Vélez-Bermúdez et al. (19).

### A Comprehensive Proteomic Inventory of pH-Responsive Proteins

The current TMT-based quantitative proteomics approach identified 40,971 and 45,844 unique peptides in shoots and roots, respectively (Fig. 4A), corresponding to 7,784 (shoots) and 14,182 (roots) proteins according to the set criteria (at least one unique peptide in two of the three biological replicates, FDR ≤ 1%). Moreover, our survey identified 7,718 and 11,257 phosphorylated peptides (with 8265 and 13,081 phosphorylation sites) from 3,633 and 10,692 phosphoproteins in shoots and roots, respectively (Fig. 4A; supplemental data set S1). The vast majority of identified phosphorylation sites were located on serine residues (79% in shoots and 86% in roots), 19% (shoots) and 13% (roots) and 2% (shoots) and 1% (roots) mapped to threonine and tyrosine, respectively (supplemental data set S1). More than a third of the identified phosphorylation sites is not listed in the PhosPhAt 4.0 data base hosted by the University of Hohenheim (https://phosphat.uni-hohenheim.de) (Fig. 4*B*), suggesting that some of the sites reported here are specific to changes in pH_*e*_. Taking the overlap between the identified proteins in shoots and roots into consideration, our study covers a total of 17,204 proteins (Fig. 4*C*), corresponding to 62% of the 27,655 protein-coding genes annotated in Araport 11 (27). A comparison of the global changes in protein abundance shows that external pH alters the proteomic profile gradually, without dramatic changes in gene expression (Fig. 4*D*-*G*).

**FIG. 4.**
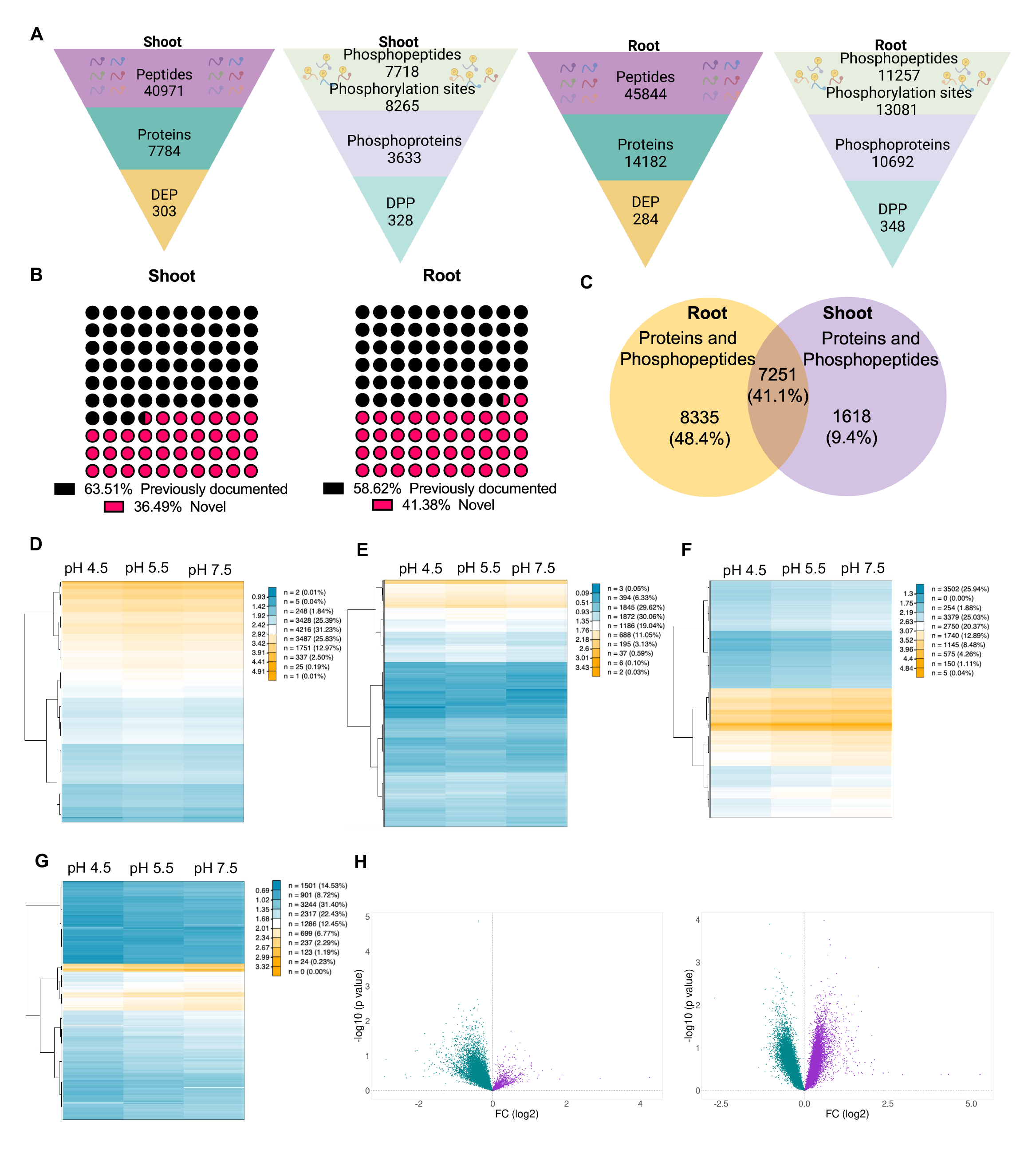
Proteomic profiling of *Arabidopsis* plants. *A*, total number of peptides, proteins, DEPs, and DPPs in shoots and roots. *B*, novel phosphorylation sites identified in this survey. *C*, overlap of identified proteins between roots and shoots. *D*, heatmap of the shoot proteome. *E*, heatmap of the shoot phosphoproteome. *F*, heatmap of the root proteome. *G*, heatmap of the root phosphoproteome. *H*, volcano plots showing up-and downregulated proteins in shoots (left) and roots (right). Upregulated proteins are depicted in purple color, cyan color indicates downregulated proteins. DEPs, differentially expressed proteins; DPPs, differentially accumulated phospho-peptides.

Generally, abundance changes were more pronounced when phosphoproteins were considered. Volcano plots revealed a bias towards decreased abundance of proteins in response to alterations in pH_*e*_ in shoots; no such bias was observed in roots (Fig. 4*H*).

Differentially expressed proteins (DEPs) between each group were defined by a fold change > 1.2 or < 0.83 and *p* < 0.05. A complete list of the DEPs is provided in supplemental data set S2. While the number of DEPs is comparable in roots and shoots, the proteins that are altered in abundance show limited concordance (Fig. 5*A*). In both pH treatments, only a small subset of proteins was differentially expressed in both organs. Generally, in pH 4.5 plants, DEPs and differentially expressed phosphopeptides (DPPs) were mostly upregulated, a response that was particularly pronounced in roots. This observation might be interpreted in terms of a more active adaptive response when plants are subjected to acidic pH relative to pH 7.5 plants (Fig. 5*B*).

**FIG. 5.**
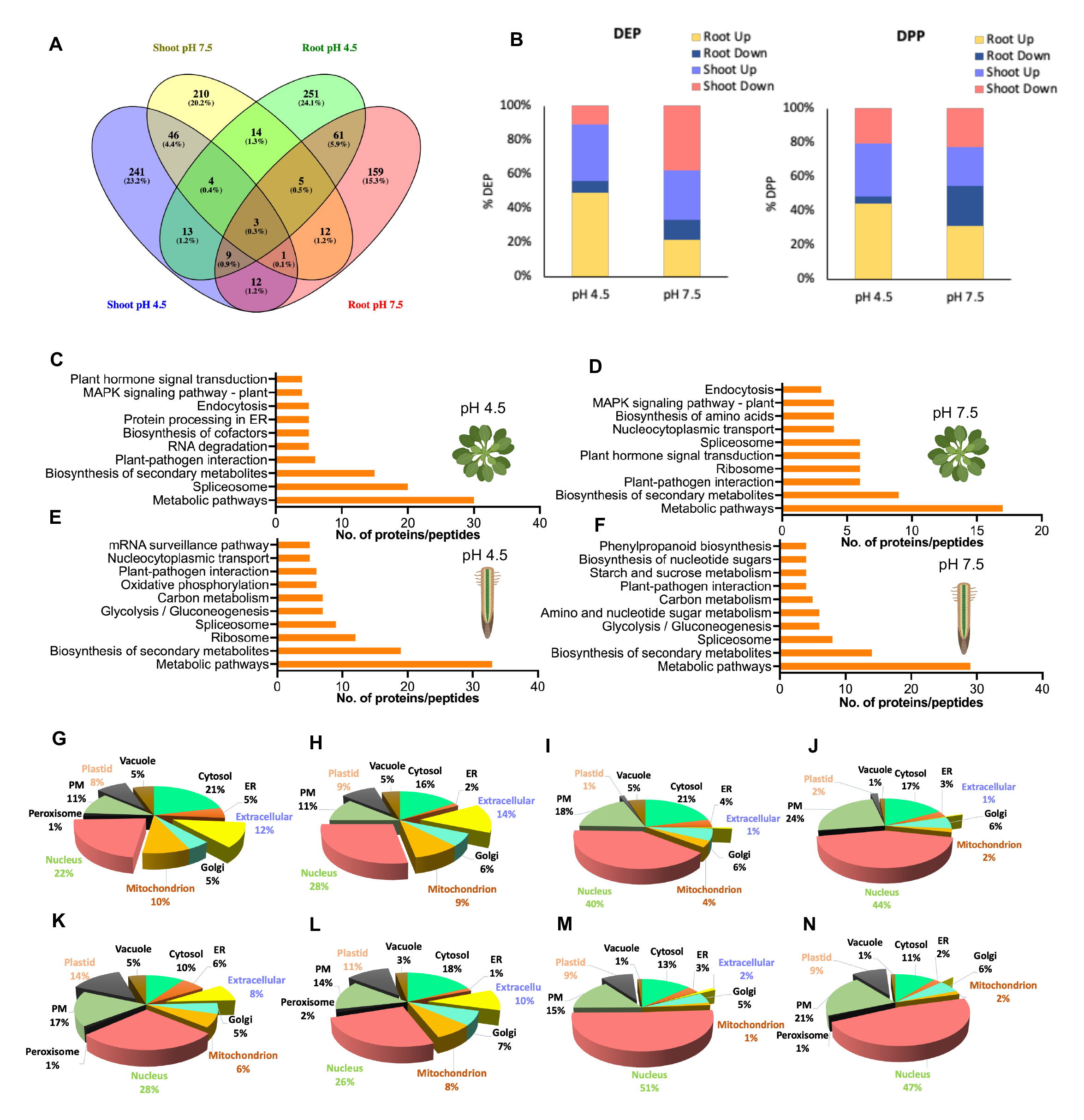
Function and localization of DEPs/DPPs. *A*, Venn diagram depicting the overlap between DEPs/DPPs among the various treatments in shoots and roots. *B*, distribution of up-and downregulated DEPs (left) and DPPs (right). *C*-*F*, KEGG analysis of DEPs/DPPs in shoots of pH 4.5 plants (*C*) shoots of pH 7.5 plants (*D*), roots of pH 4.5 plants (*E*), and roots of pH 7.5 plants (*F*). *G-N*, subcellular localization of DEPs/DPPs in roots of pH 4.5 plants (*G*), DEPs in roots of pH 7.5 plants (*I*), DPPs in roots of pH 4.5 plants (*J*), DPPs in roots of pH 7.5 plants (*K*), DEPs in shoots of pH 4.5 plants (*L*), DEPs in shoots of pH 7.5 plants (*M*), DPPs in shoots of pH 4.5 plants (*N*), DPPs in shoots of pH 7.5 plants (*O*).

KEGG pathway analysis revealed that in shoots of pH 7.5 plants the category ‘biosynthesis of secondary metabolites’ was most overrepresented, followed by ‘plant-pathogen interaction’, ‘ribosome biogenesis’, and ‘plant hormone signal transduction’ (Fig. 5*D*). In shoots of pH 4.5 plants, splicing-related processes were most represented, followed by ‘biosynthesis of secondary metabolites’ and ‘plant pathogen interaction’ (Fig. 5*C*). In roots, the categories ‘biosynthesis of secondary metabolites’, ‘splicing’, and ‘carbon metabolism’ were enriched; no pronounced differences were observed between the two pH values under study (Fig. 5*E*, *F*).

A notable difference between the identified DEPs and DPPs is related to the subcellular localization of the proteins. While – independent on the organ or whether acidic or alkaline pH is considered – a substantial portion of DEPs was located in the extracellular space, this fraction was very small (1-2%) of the total DPPs (Fig. 5). A similar shift was observed for DEPs and DPPs localized to mitochondria. This change in ratio was inverse between the two fractions when nuclear-located proteins were considered. Here, mostly phosphorylated peptides were differentially enriched, indicative of possible transcriptional control by protein phosphorylation.

Moreover, when compared to alkaline pH, DEPs localized to the ER were more abundant in acidic conditions, indicating pH-dependent changes in the ERs function.

### pHe Governs Transport Processes Across the Plasma Membrane

The DEPs and DPPs identified here comprise a suite of plasma membrane-bound transporters, suggesting that the import and export of various substrates is governed by pH_*e*_. In roots, in this population changes in DPPs were more pronounced than alterations in protein abundance, indicative of chiefly post-translational regulation of plasma membrane-localized proteins. Exposure to low pH increased the expression of a peptide corresponding to the Rapid Alkalization Factor proteins RALFL22, RALF23, and RALFL33. RALF peptides alkalize the apoplast by inhibiting H^+^-ATPases on the plasma membrane through the receptor FERONIA (28). In the present study, AHA1 peptides phosphorylated at S931 increased upon exposure to low pH, presumably leading to reduced efflux of protons and alkalization of the apoplast. Since a large portion of the phosphorylation sites identified here are uncharacterized, our predictions regarding activation/repression are made based on the structure, physiological role, and closeness to previously described sites (supplemental Table S1; Fig. S1).

The expression of the polyamine transporter ABCG28 was increased upon exposure to acidic conditions (Fig. 6*A*). Another ABC-type transporter, PDR7, showed increased phosphorylation at S824 in response to low pH, which may lead to an increase in activity and, possibly, to a change in substrate specificity (Fig. 6*A*). PROT2, a plasma membrane-bound transporter with affinity to betaine, proline, and GABA, was downregulated at low pH (Fig. 6). In PDR8, a close homolog of PDR7 (83% identity), phosphorylation represses the transport of indole-3-butyric acid (IBA) on the expense of camalexin, thereby prioritizing defense over growth. Here, S823 and S825 serves as phospho-switches (29). A similar phospho-switch may alter substrate specificity of PDR7, prioritizing defense over growth at low pH. At high pH, a decrease in phosphorylation was observed for PDR7 (at S43) and for PDR8 (at S45), a modification that possibly supports IBA transport in alkaline conditions (supplemental Fig. S2). Also the auxin effluxer ABCB15 (MDR13) (30) showed decreased phosphorylation in response to high pH (Fig. 6*A*).

**FIG. 6.**
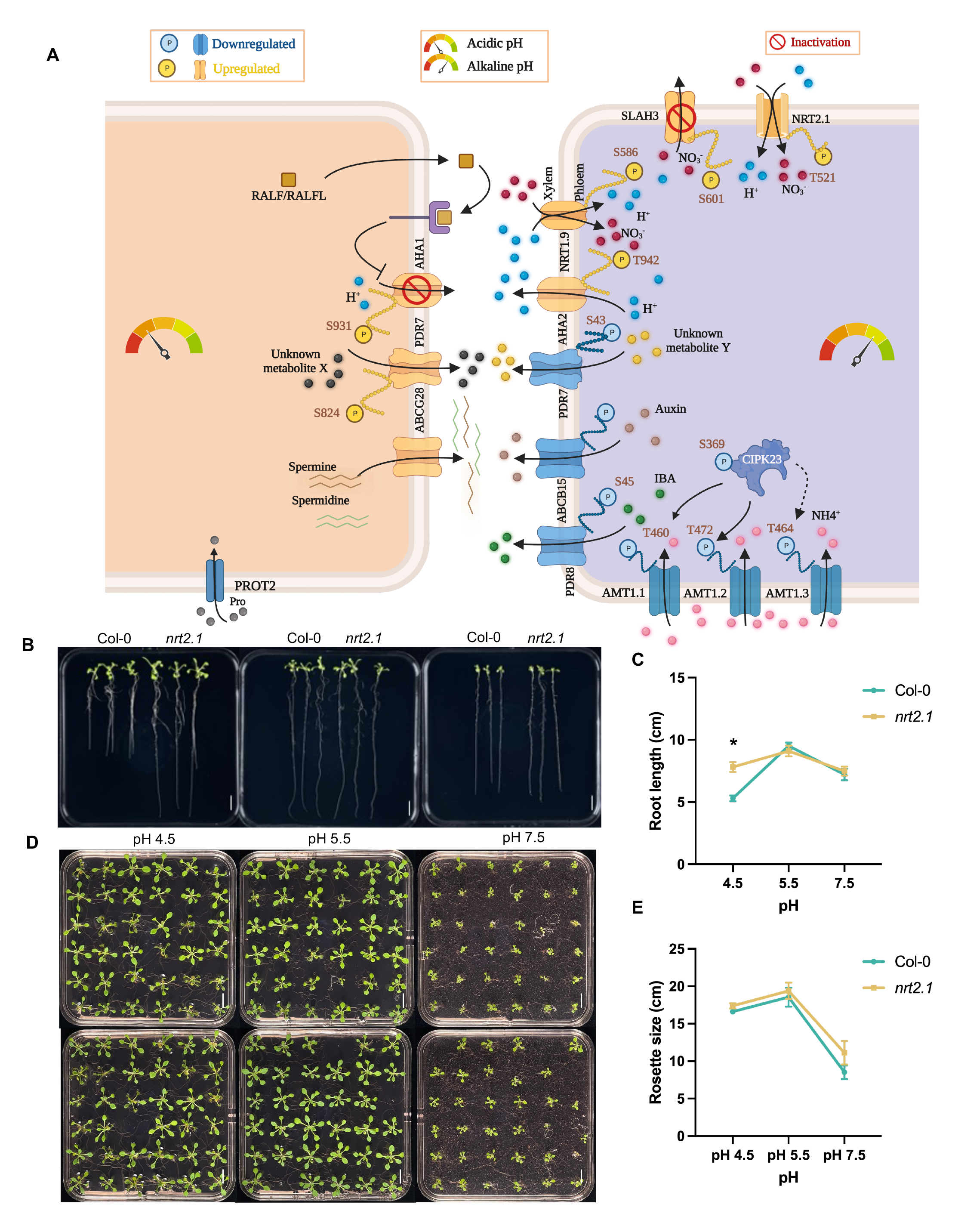
pH_*e*_-responsive transmembrane transport processes. *A*, pH-dependent regulation of plasma membrane-localized transporters. Transporters that change in abundance or are differentially phosphorylated upon exposure to acidic (left) or alkaline (right) media are depicted. *B*, root phenotype Col-0 and *nrt2.1* grown for 14 days on media adjusted to pH 4.5, pH 5.5, or pH 7.5. *C*, two-way ANOVA was performed to detect significant differences in root length between the genotype under variable pH conditions. Black asterisks represent significant differences between Col-0 and *nrt2.1* according to analysis of variance, Tukey’s test (*P* < 0.05). *D*, Rosette phenotype of wild-type and *nrt2.1* plants. *E*, two-way ANOVA test for significant differences in rosette size between wild-type and *nrt2.1* mutant plants. Scale bar = 1 cm. Error bars = SEM. *n* = 5 biologically independent samples per genotype.

### Alkaline pHe Massively Induces Trans-Plasma Membrane Nitrogen Transport

In pH 7.5 plants, AHA2, the major P-type ATPase isoform on the plasma membrane besides AHA1, exhibited increased phosphorylation at T942, a site associated with activation of the enzyme (supplemental Table S1). Moreover, exposure to alkaline media enhanced the expression and altered the phosphorylation states of both nitrate and ammonium transporters, suggesting massively increased import of nitrogen. This finding was unexpected, since plants were grown at a nitrate concentration that allows luxury (i.e., more than adequate) consumption of nitrogen. Phosphopeptides associated with the plasma membrane-localized nitrate transporters NRT2.1 (high affinity) and NRT1.9 (low affinity) accumulated at high pH (Fig. 6*A*). NRT2.1 was phosphorylated at T521, NRT1.9 at S586; both sites are likely to activate the transporters (supplemental Table S1, Fig. S1). While NRT2.1 is involved in the acquisition of nitrate from the soil solution, NRT1.9 is expressed in companion cells in the root phloem and is mediating the loading of nitrate into the phloem (31, 32). Notably, NRT1.9 facilitates the downward transport of nitrate and enhance the uptake of nitrate into root cells (32).

The S-type nitrate channel SLAH3 was phosphorylated at S601 upon exposure to alkaline pH (Fig. 6*A*). Phosphorylation at this site is mediated by the SNF1-related protein kinase SnRK1.1, inhibiting SLAH3 activity to prevent nitrate loss (33). SLAH3 can sense cytosolic acidosis, which activates the channel by monomerizing the (default) SLAH3 dimers via protonation of histidine 330 and 454 (34). Our data show that exposure to alkaline media - likely coupled to a slight increase in pH_*cyt*_ - deactivates the channel via phosphorylation and, possibly, by deprotonation of H330 and H454.

Phosphopeptides of three ammonium transporters from the AMT family decreased in abundance upon exposure to alkaline media (Fig. 6*A*). Decreased phosphorylation was observed at T460 (AMT1;1), T472 (AMT1;2), and T464 (AMT1;3), which likely derepressed the channels (supplemental Table S1). Phosphorylation of ATM1;1 and AMT1;2 is mediated by CBL-INTERACTING PROTEIN KINASE 23 (CIPK23) (35). In pH 7.5 plants, CIPK23 showed decreased phosphorylation, a modification that decreased the activity of the kinase (Fig. 6*A*). Thus, alkaline conditions appear to increase the import of ammonium via AMT channels.

To further investigate a putative role of nitrogen transport in pH homeostasis, we focused on the transporter NRT2.1. In addition to altered phosphorylation, *NRT2.1* was found to be upregulated in response to alkaline media pH in a previously conducted transcriptomic survey (12). To identify putative pH-dependent phenotypes of *nrt2.1*, we grew homozygous mutant plants on media adjusted to the pH values under study. The mutant allele was characterized previously (36). While the growth pattern of the mutant did not deviate from the wild type regarding rosette size and root length when grown at pH 5.5 or 7.5, the root growth reduction typically observed in wild-type plants at pH 4.5 was not apparent in *nrt2.1* mutant plants, suggesting a moonlighting role of NRT2.1 in pH-dependent root growth regulation (Fig. 6).

### pHe Modulates Sugar Transport and Metal Homeostasis

pH-induced changes in metal and sugar transport are depicted in Figure 7. Exposure to low pH caused increased phosphorylation of the vacuolar iron transporter VTL5 (Fig. 7). While it has not yet been established whether phosphorylation at this site activates or deactivates the transporter, it can be assumed that iron sequestration is increased at low pH in anticipation of the increased availability of the nutrient associated with acidic conditions. Since in the present study iron is provided as pH-stable FeEDDHA and pFe was supposedly unchanged over the pH values under study, altered abundance and phosphorylation of VTL5 does not appear to reflect a response to altered iron uptake.

**FIG. 7.**
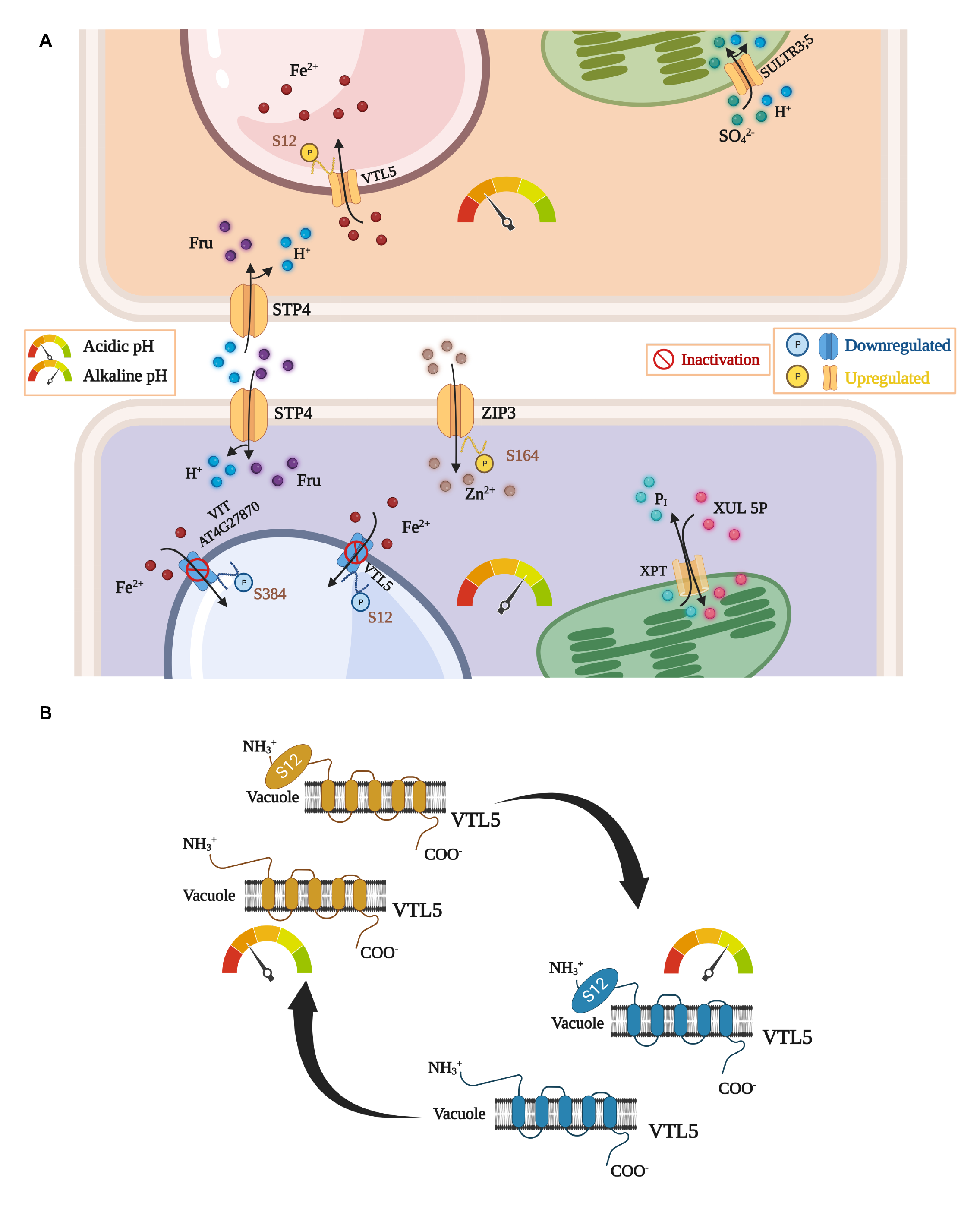
pH-responsive metal and sugar transporters in roots. *A*, regulation and intracellular localization of pH-dependent transporters in response to exposure to acidic (upper panel) and alkaline pH (lower panel). *B*, regulatory switch of VTL5 in response to changes in pH_*e*_ via alterations in protein abundance and phosphorylation at S12.

Furthermore, the sulfate transporter SULTR3.5, residing in the plastid inner envelope membrane, and the plasma membrane-bound sugar/H^+^ symporter STP4 that provides plastids with xylose-5-phosphate, showed increased expression under acidic conditions (Fig. 7). SULTR3.5 may be employed to reduce cytosolic acidosis by sequestering cytosolic protons in the plastid via SO ^-^/H^+^ co-transport. Under alkaline conditions, STP4 was induced together with the zinc transporter ZIP3, which is phosphorylated at S164. STP4 protein abundance was previously shown to be responsive to the iron status of the plants (37). In contrast to pH 4.5 plants, alkalinity repressed phosphorylation of VTL5 (Fig. 7). Moreover, phosphopeptides of another vacuolar iron transporter, VIT, also showed reduced abundance, indicative of repressed cellular iron sequestration (and anticipated reduced availability of iron) under these conditions. Thus, sequestration of iron into the vacuole via VTL5 is regulated in a pH-dependent manner, both by accumulation of protein and increased phosphorylation at low pH (Fig. 7*B*). Of note, *VTL5* was also found to be responsive to the iron status and to pH_*e*_ at the transcriptional level (12, 38).

### Root Responses to pHe Are Partly Conserved in Shoots

In shoots, low pH activated AHA2 through induced phosphorylation at T980 (Fig. 8). Similar to what has been observed in roots, abundance of the sucrose transporter STP4 was increased in leaf cells of pH 4.5 plants. Furthermore, the export of copper from the vacuole was repressed through decreased abundance of the copper transporter COPT5. In pH 7.5 plants, the low-affinity nitrate transporter NRT1.12 showed increased expression, resembling the promotive effect of alkalinity on NO_3_ ^-^/H^+^ transport in root cells. NRT1.12 is predominantly expressed in companion cells of the major vein and putatively involved in the redistribution of nitrate derived from the xylem (39). Similar to roots, reduced phosphorylation at T460 was observed for AMT1.1 in shoots upon exposure to high pH, indicative of increased ammonium uptake. Furthermore, PDR8 phosphorylation was decreased in leaves of pH 7.5 plants. These data suggest that leaf cells respond vividly to changes in pH_*e*_, with a subset of the responses being conserved between the two organs.

**FIG.8. pH-dependent.**
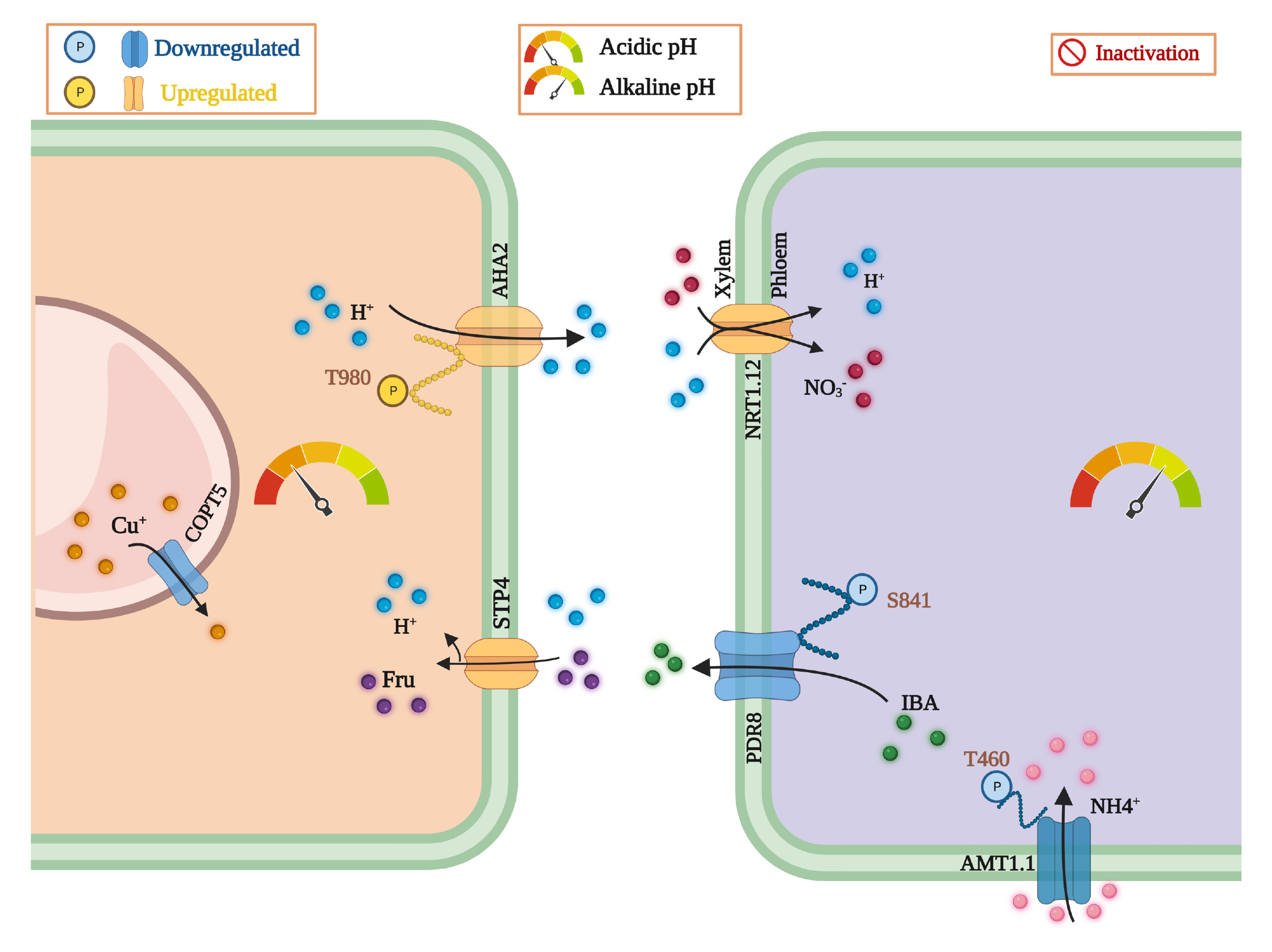
transporters in the shoot. Regulation and intracellular localization of pH- dependent transporters in response to exposure to acidic pH (left panel) and alkaline media (right panel).

### A Subset of Genes is Responsive to pHe Both on Protein and Transcript Levels

The DEPs identified here showed a substantial overlap to a previously published data set derived from roots of plants subjected to short-term exposure to pH 7.5 (12). This suite of 66 genes includes NRT2.1, AMT1.1, VTL5, CIPK23, PDR8, STP4, and ABCB15, indicating that some of the processes described in the present study are subject to robust regulation, modulating the steady-state abundance of both transcripts and proteins (supplemental Table S2). Interestingly, ABCB15 is co-expressed with several genes identified in the current proteomics survey (i.e., *NRT1.9*, *AMT1.3, AHA2*), indicative of a module of genes that alters fluxes across the plasma membrane in response to alkaline conditions. The most downregulated gene in both surveys was the glutamine amidotransferase-like superfamily protein At5g38200.

Similarly, a suite of DEPs of plants grown at low pH was previously described as being responsive to short-term exposure to low pH at the transcript level (11). DEGs that were identified in both surveys included, among others, the auxin transporter PILS5, the DUF642 protein DGR2, the polygalacturonase-inhibiting protein PGIP1, and EXORDIUM LIKE 2 (EXL2). PILS5 is an ER-localized auxin transporter critical to cellular auxin homeostasis (40). In pH 4.5 plants, PILS5 was upregulated both at the transcript and phospho-protein level. EXL2, localized to the extracellular space, was found to be responsive to hypoxia, which causes a reduction in pH_*cyt*_ (41). EXL2 is co-expressed with *RALF33*, *PGIP1*, and *STOP1*, and was shown to be induced by exposure to low pH at the transcript (11) and protein level (this study). Notably, *PILS5*, *DGR2*, *PGIP1*, and *EXL2* are putative targets of STOP1 (42). Taken together, this comparison suggests that the current data set is consistent with the expected inverse regulation of congruent transcripts-protein pairs in response to high and low pH, thus validating the proteomic approach.

## DISCUSSION

*Unfavorable pH*e *Induces pH-Specific Growth Patterns*

Growth of (mildly calcifuge) *Arabidopsis* plants was optimal at moderately acidic pH_*e*_ and markedly impaired when the media pH was lowered to 4.5 or increased to 7.5. Growth cessation occurred, however, in a pH-specific pattern. Primary root growth was much more restricted at pH 4.5 when compared to alkaline conditions, possibly to prevent excess accumulation of Al^3+^, NH_4_ ^+^, and Mn^2+^ ions that are highly abundant in acid soils at pH values below 5. Notably, rosette size was not significantly reduced under acidic conditions, suggesting that low pH specifically restricts root growth to avoid rhizotoxicity. This scenario is not likely to account for the growth reduction observed at high pH. Rather, the low proton concentration in the apoplast counteracts cell wall loosening and restricts cell expansion as postulated by the acid growth theory (4, 43). In contrast to low pH plants, exposure to pH 7.5 media severely reduced rosette size, suggesting that suboptimal performance at alkaline pH is adopted as a means to decrease the demand for mineral nutrients that are lowly abundant under such conditions, i.e., phosphate, zinc, and iron.

Alkaline conditions not only restricted growth, but also severely compromised the fitness of the plants. Our analysis shows that flower morphology was altered in a way that restrict self-pollination and massively reduced the seed number when plants were grown in alkaline soil. Moreover, the number of pollen grains produced per plant was greatly diminished. These observations suggest that *Arabidopsis thaliana*, at least the Col-0 accession, is not adapted to thrive in even mildly alkaline soils. Thus, rhizo-toxicity on one end and reduced fitness on the other end of the pH scale investigated here limit the ecological amplitude of *Arabidopsis thaliana*.

### A ‘Reduced Bias’ Approach to Concurrently Map the Proteome and Phosphoproteome

*In situ*, rapid changes in soil pH necessitate strategies to maximize growth whenever the proton concentration in the rhizosphere deviates from a ‘just right’ scenario. The proteomic survey reported here - conducted to infer such strategies - identified a total of 17,204 proteins in roots and shoots of *Arabidopsis* seedlings, corresponding to approximately 62% of the protein-coding genes. This number is close to the recent draft of the *Arabidopsis thaliana* proteome (44) and allows for a coverage that is still somewhat lower but close to the comprehensiveness of transcriptomic analyses such as RNA-seq.

The overlap between the current proteomic study and previously conducted transcriptomic profiling in response to short-term exposure to either alkaline (pH 7.5 (12)) or acidic (pH 4.5 (11)) media was moderate, indicative of extensive regulation at the post-transcriptional level. This observation is in line with a generally low concordance of protein and mRNA expression in plants (13). However, several key processes appear to be conserved at both levels, indicating that expression changes of a suite of genes mediating critical adaptations to pH_*e*_ result in robust alterations in the steady-state levels of their cognate proteins. The congruent subset of genes/proteins comprised the uptake of nitrogen (via NRT2.1 and AMT1.2) and the export of auxin (via ABCB150 and PDR8) under alkaline conditions as well as auxin homeostasis (PILS), defense (PGIP1, DRG2), and pH homeostasis (EXL2) under acidic conditions. Moreover, our study shows that changes in pH_*e*_ lead to extensive alterations in protein phosphorylation, indicating that a large part of the acclimation to pH_*e*_ depends on post-translational signaling events. A large subset of the circa 43,000 phosphosites identified here are not listed in comprehensive data bases, suggesting that some - or even most - of these so far undocumented events are specific to changes in pH_*e*_.

### Proton Transport Across the Plasma Membrane Recalibrates Cellular pH Homeostasis

Two auxin-triggered mechanisms, i.e., H^+^ export through activation of plasma membrane proton pumps and net cellular H^+^ influx modulated by TIR1/AFB signaling, constitute a ‘gas and brake’ mechanism that govern root growth (43, 45). While, generally, low apoplastic pH supports cell expansion, a too acidic environment can arrest growth and compromise nutrient acquisition (46). Under acidic conditions, we observed increased abundance of (one or more) RALFs, cysteine-rich peptides that counteract growth by inhibiting ATPase-mediated H^+^ extrusion (47). RALF33 was shown to inhibit AHA activity through a signaling cascade that involves cytosolic Ca^2+^ signatures, possibly by phosphorylation of S931 through PKS5 (28, 48). Extracellular alkalinization by RALF peptides is mediated through several processes that may include various receptors (28). The RALF peptide identified here might account for the decreased activity of AHA1 (which was phosphorylated at S931) in pH 4.5 plants. By contrast, in pH 7.5 plants, AHA2 was activated via phosphorylation at T942. While this site is so far uncharacterized, phosphorylation of a very close serine residue (S944) was shown to lead to increased H^+^ efflux (supplemental Table S1; Fig. S1). Thus, the dominating (early) electrogenic H^+^ movements in response to alterations in pH_*e*_ are 1) a decrease in H^+^ efflux in low pH plants through reduced activity of AHA1, possibly to limit acidification of the apoplast, and 2) an increase in H^+^ efflux via AHA2 in high pH plants, likely to support anion/H^+^ symport processes.

The concurrent induction of a suite of nitrogen transporters in response to alkaline conditions was a surprising observation. However, at least for NRT2 transporters this finding was not completely unexpected. Increased expression of nitrate transporters, including *NRT2.1*, was reported in a transcriptional survey of plants subjected to short-term exposure to high pH (12). Accumulation of an NRT2.1 homolog in response to high pH was also observed in a proteomic study on sugar beet plants (49), suggesting that this response is potentially conserved across species. The exact role of NRT2.1 in pH homeostasis remains to be established. A previous report showed that in Arabidopsis *NRT2.1* was strongly downregulated at low pH, while the dual-affinity transporter *NRT1.1* was upregulated in a STOP1-dependent manner under these conditions (50). Both transporters are nitrate-inducible (51). The inverse response of *NRT1.1* and *NRT2.1* to pH_*e*_ suggests that distinct NRTs are employed for distinct tasks. In contrast to *nrt2.1* mutant plants, in which no significant root growth cessation was noted at low pH, both *nrt1.1* and *stop1* mutants showed a more pronounced root growth inhibition than the wild type under such conditions (50). Moreover, treatments compromising root meristem activity such as excess iron, toxic levels of Al^3+^ ions, low pH, and low phosphate availability decreased *NRT2.1* transcript levels (42), supporting a role for NRT2.1 in repressing root growth. NRT2.1 was shown to negatively regulate lateral root formation (52), supporting this supposition. Of note, *NRT2.1* was within the small suite of genes that is regulated transcriptionally (12), post-transcriptionally, and post-translationally (by phosphorylation), indicating robust multilevel regulation of this transporter. In addition to increased nitrate uptake, three AMT-type ammonium transporters, mediating the uptake of ammonium from the soil solution, were upregulated in pH 7.5 plants via decreased (repressive) phosphorylation. Thus, the uptake of nitrogen appears to be generally supported under alkaline conditions.

The reasons as to why nitrogen influx is increased in response to high pH remain to be elucidated. All changes in nitrogen transport observed in pH 7.5 plants potentially aid in acidifying the cytoplasm. Alkaline pH_*e*_ creates a H^+^ gradient from the cytoplasm to the apoplast, which supports the efflux of H^+^ ions and gradually alkalizes the cytoplasm. Nitrate uptake is associated with cytosolic acidosis due to the concomitant transport of two protons per nitrate molecule across the plasma membrane (53-55). In contrast to NRT2.1, AMTs are NH_4_ ^+^ uniporters (56). Thus, NH_4_ ^+^ transport is electrogenic with most of the NH4^+^ remaining undissociated after crossing the membrane (57). However, a small fraction of the influxed NH_4_ ^+^ dissociates into NH3 and H^+^ and acidifies the cytosol (58). Given that the concentration of free cytosolic H^+^ is in the sub-micromolar range (59), this process will significantly affect pH_*cyt*_. The acidosis associated with the uptake of protons via NO_3_ ^-^-H^+^ co-transport is thought be compensated by the (H^+^-consuming) assimilation of nitrate (60). Conspicuously, the activation of nitrogen transporters in response to elevated pH_*e*_ does not seem to lead to increased assimilation of the nutrient. This observation stands in contrast to nitrate-treated plants in which induction of nitrate transporters is coupled with increased expression of genes coding for enzymes involved in the formation of organic nitrogen compounds (51). These observations suggest that the object of desire here is the change in cytosolic H^+^ concentration rather than the nitrogen as such.

### pHe Governs Transition Metal Homeostasis

The availability of iron is intricately linked to soil pH, causing an approximately one-thousand-fold decrease for each one-digit increase in pH (61). This does, however, not apply to the current experimental setup in which iron is provided in a highly stable form, which makes it unlikely that the iron status of the plant differs among the treatments. Only two proteins of the Arabidopsis ‘ferrome’ (a suite of genes the expression of which is robustly responsive to iron (62)), i.e., the plasma membrane-bound zinc transporter ZIP3 and the vacuolar iron transporter VTL5, are responsive to pH_*e*_. The activity of VTL5, a member of a small family of nodulin-like genes with homology to AtVIT1 and ScCCC1p (63, 64), is regulated by both expression and phosphorylation and is one of the few proteins that are inversely regulated in response to acidic and alkaline media pH. In addition, we observed decreased phosphorylation of the VIT family protein At4g27870 in response to high pH, putatively downregulating vacuolar iron sequestration. While iron transport activity for At4g27870 remains to be experimentally verified, the response of both vacuolar transporters seems to reflect rather an anticipation of iron shortage generally associated with alkaline pH than an actual change in cytosolic iron level. From the vast suite of responses to alterations in iron availability, vacuolar iron sequestration appears to be the only process that is responsive to pH_*e*_, suggesting that control of the cytosolic iron concentration has priority over mobilization, uptake, and xylem-loading of iron (to ultimately satisfy the demand of sink tissues). Thus, avoidance of iron overload - strictly controlled by pH_*e*_ – seems to be controlled separately from the responses to low iron availability, which induces a much more pronounced and multi-faceted response (65).

While we observed promotive phosphorylation of the Zn transporter ZIP3 in response to high pH (and, supposedly, reduced iron availability), Zn^2+^ uptake was found to be downregulated upon iron deficiency to avoid accumulation of excess Zn^2+^ through the promiscuous Fe^2+^ transporter IRT1, which is induced upon iron deficiency (66). These seemingly contrarily observations suggest that phosphorylation at high pH occurs in anticipation of the decreased phyto-availability of zinc generally associated with such conditions, a response that might interfere with or be overruled by transcriptional repression caused by iron deficiency. It may thus be assumed that the activity of metal transporters is orchestrated by distinct signaling cascades to optimally acclimate transport activities to the prevailing conditions. These data suggest that pH_*e*_ is controlling a distinct subset of processes that is independent of the responses to the (external) availability of the metals or the (internal) status of the plant.

### ABC Transporters Govern Growth-Defense Trade-Offs and Tune pHapo

Several ABC-type transporters were differentially expressed in response to pH_*e*_, controlled by both protein abundance and phosphorylation. Post-translational modifications of this type of transporters not only change their activity, but may also have consequences for their substrate specificity. For instance, PDR8 prioritizes either IBA or camalexin transport, a discretion governed by the phosphorylation state of the transporter (29). In the case of PDR8, phosphorylation - mediated by the LRR receptor-like kinase KIN7 - decreases the efflux of the ‘canonical’ substrate IBA and, as a consequence, favors the export of the phytoalexin camalexin (29). The phosphorylation sites of PDR8 are arranged in two clusters (cluster 1: S40 and S45; cluster 2: S823 and S825) preceding the nucleotide binding domains (29). Phosphorylation of PDR8 at S45 decreased in pH 7.5 plants, indicating that IBA export and, thus, growth is favored under these conditions. Upon exposure to acidic conditions, PDR7 - the closest homolog of PDR8 - was phosphorylated at S824 (cluster 2, supplemental Fig. S2). Similar to what was observed for PDR8, phosphorylation of PDR7 was repressed when plants were subjected to alkaline conditions, leading to reduced abundance of peptides that are phosphorylated at S43 (cluster 1). Assuming functional redundancy between the two transporters, the question whether to grow of fight appears to be decided by a pH-dependent phospho-switch of the transporters. In this scenario, export of IBA would be repressed at low pH on the expense of camalexin while at high pH IBA export is prioritized.

ABCB15 is predominantly expressed in the vasculature and was functionally associated with auxin transport (30). In the present study, phosphorylation of ABCB15 was decreased at alkaline pH. While it is unclear at present whether this modification decreases or increases the activity of the transporter, the former case is more likely since transcript abundance of this gene was strongly reduced upon exposure to high pH and by supplementing the growth media with bicarbonate (12, 67). The pH-dependent regulation of ABCB15 suggests a possible involvement of this transporter in the regulation of auxin-induced cell elongation.

A different role may be attributed to ABCG28. The transporter was described as being preferentially expressed in pollen grains and pollen tubes, where it contributes to the apical accumulation of ROS (68). In the present study, ABCG28 protein accumulated in roots of plants grown at low pH. *ABCG28* is not transcriptionally regulated by pH (11), suggesting that the gene is mainly or entirely post-transcriptionally regulated. ABCG28 transports polyamines, which – in addition to a range of other functions - were suggested to play a role in the regulation of pH_*cyt*_ (69). Ectopic expression of *ABCG28* in root hairs led to improved root hair growth at high pH_*e*_, likely due to the secretion of polyamines (68). The function of ABCG28 in roots in pH homeostasis is still elusive. In bacteria, the polyamine putrescine neutralizes an acidic external environment and increase survival in extreme acids (70). This response is similar to the production of the alkaline compound cadaverine in response to low external pH *E. coli* via the CadC-mediated signalling system (71). Here, cytoplasmic pH is increased by cadaverine produced via cytoplasmic lysine decarboxylase. Cadaverine is exported to the periplasmic by a membrane-integrated lysine/cadaverine antiporter to increase the extracellular pH. While these observations do not allow to infer the precise function of the transporter, it tempting to speculate that ABCG28 is employed at low pH to tune pH_*apo*_ by secreting polyamines into the extracellular space, a supposition that is supported by the predominant expression of ABCG28 in the outer cell layers of the root (http://bar.utoronto.ca).

### Leave Cells Respond Competently to Rhizospheric pH

Albeit pH_*e*_ is a cue that is not expected to directly affect the function or growth of above-ground tissues, the response of roots and above-ground plant parts was roughly similar in robustness when the number of differentially expressed proteins is taken as a criterium. While the physiological processes that are modulated by pH_*e*_ differed substantially between roots and shoots, some responses such as the induction of NO_3_ ^-^-H^+^ co-transport, NH4 uptake via AMT1.1 as well as IBA export via PDR8 were conserved between roots and shoots, in particular in pH 7.5 plants. Together, these observations suggest that changes in pH_*e*_ perceived by rhizodermal cells can be relayed to shoots.

Edaphic cues such as salt stress or drought perceived by roots were shown to trigger a transient increase in leaf apoplastic pH (72), corroborating the idea of inter-organ propagation of abiotic and, possibly, biotic stimuli. Transitory changes in pH_*apo*_ were recognized as possible ‘general stress factors’, systemically reporting edaphic information (73). Such inter-organ communication could be mediated by Ca^2+^ ions. Interestingly, Ca^2+^ transients in the cytosol are accompanied by alterations in pH, suggesting that integrated Ca^2+^/H^+^ signaling can orchestrate downstream events (74). Calcium waves can migrate through the plant (75, 76) and might be employed to convey information on pH_*e*_ to above-ground plant parts. It remains to be shown, however, how such Ca^2+^/H^+^ signals are translated into cellular processes that ultimately dictate the physiological readouts triggered by pH_*e*_. Candidates for decoding such information are calcium-binding proteins and calcium-dependent protein kinases (CPKs), a cluster of which was found to be differentially phosphorylated in response to changes in pH_*e*_ in both roots and shoots (Fig. 9). CPKs were shown to be involved in cell expansion and plant immunity (77) and may serve as nodes to gather and translate information on the H^+^ concentration into growth and immune responses. Besides CPKs, the group of proteins related to calcium signaling comprises several CaLB-domain proteins and calcium-binding EF hand proteins, which are chiefly post-translationally regulated. Also, the Na^+^-Ca^2+^ exchanger NCL, previously shown to regulate auxin responses (78), was responsive to the pH regime. Notably, in pH 4.5 plants, alterations in phosphorylation were most pronounced in roots, while in pH 7.5 plants such changes were more prominent in shoots, with phosphorylation events increased in the former and decreased in the latter case (Fig. 9). These observations are indicative of intensive whole plant calcium signaling triggered by changes in pH_*e*_.

**FIG. 9.**
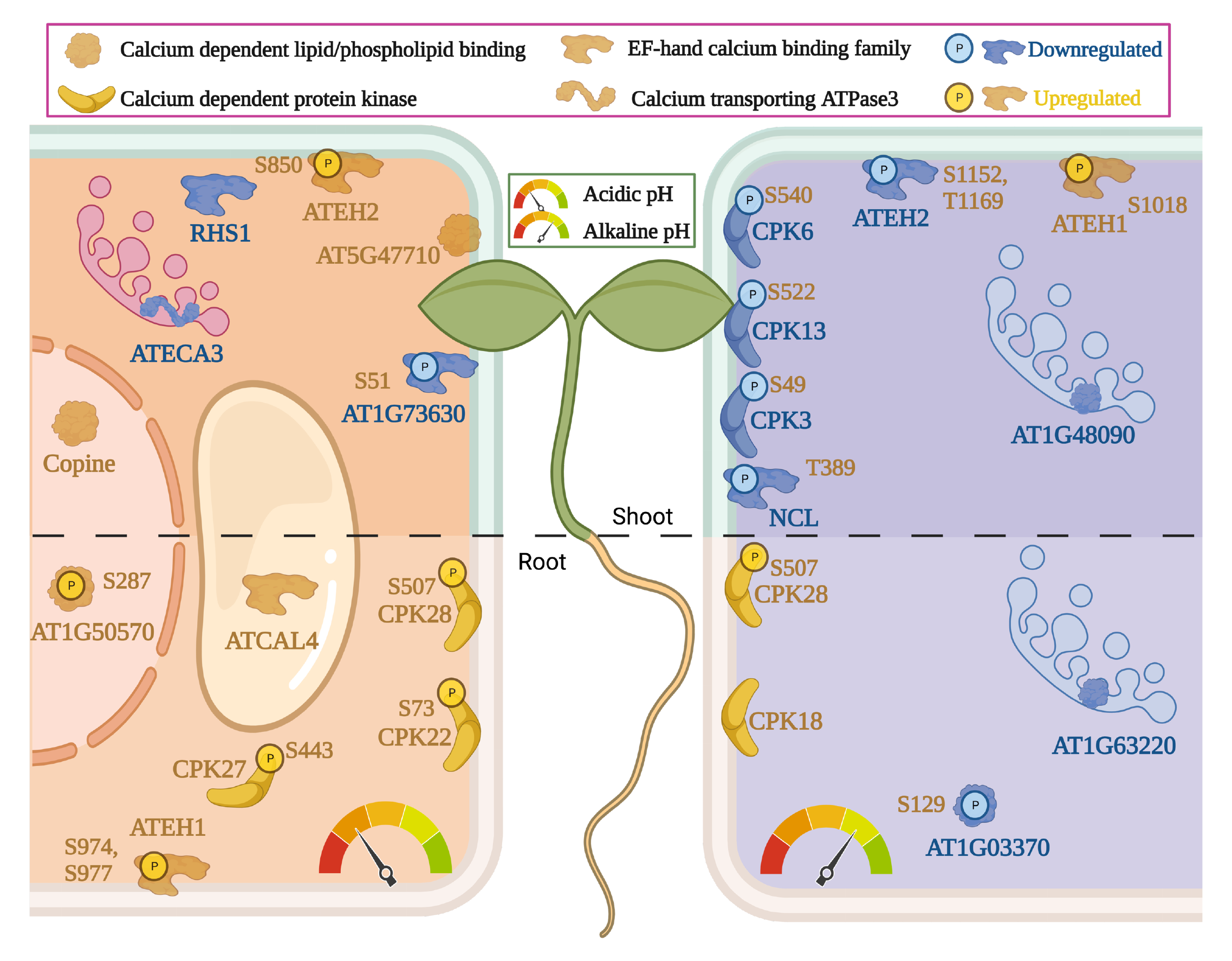
Calcium-responsive DEPs in shoots and roots. Regulation and intracellular localization of pH-dependent transporters in response to exposure to acidic pH (left panel) and alkaline media (right panel).

Pronounced pH_*e*_-induced changes in phosphorylation pattern were also observed in the (Ca^2+^-binding) plasma membrane-localized proteins AtEH1/Pan1 and AtEH2/Pan1, components of the TPLATE complex (TPC) (79). TPC is controlling growth and perception of environmental signals via its role in clathrin-mediated endocytosis (80). TPC may also be involved in the regulation of H^+^-fluxes across the plasma membrane. A mutant harboring defects in the large TPLATE subunit TASH3 – referred to as *nosh* (no SH3) - showed reduced internalization of TRANSMEMBRANE KINASE 1 (TMK1) (81). TMK1 is required for auxin-induced activation of plasma membrane H^+^-ATPases via phosphorylation, promoting cell wall acidification and cell elongation (82). Homozygous *nosh* seedlings exhibited smaller rosette leaves and reduced root and hypocotyl length relative to wild-type plants (81). Similar to what was observed here for plants grown in alkaline soil, *tash* mutant plants showed reduction in the viable pollen number. In yeast, phosphorylation of the AtEH1/AtEH2 homolog Pan1p inhibits endocytic functions (83, 84). In plants, the mechanisms and consequences of AtEH1/AtEH2 phosphorylation have not yet been investigated. Our study shows that alterations in pH_*e*_ result in differential phosphorylation of AtEH1 and AtEH2 at various sites in both leaves and roots, suggesting that pH-dependent changes in AtEH1/AtEH2 activity participate in the acclimation of growth to the prevailing environmental conditions. Of note, only one (S974) of the various phosphorylation sites of AtEH1 and AtEH2 reported here has been previously annotated, suggesting the employment of specific kinases to regulate clathrin-mediated endocytosis in a pH_*e*_-dependent manner.

## CONCLUSIONS

Our data show that – similar to the pronounced changes observed previously at the transcriptional level – pH_*e*_ modulates the abundance of a large subset of proteins involved in a variety of processes aimed at adjusting growth and tuning pH_*cyt*_ and pH_*apo*_. Such changes were particularly pronounced for phosphopeptides, indicative of intensive signaling triggered by short-term changes in media pH. It can be further deduced from the data that the regulation of growth, the balance between growth and defense, and the transport of H^+^ ions across the plasma membrane are the central themes associated with the recalibration of pH homeostasis. Nitrate-H^+^ cotransporters seem to play specific roles in the adaptation of plants to pH_*e*_. The multi-faceted regulation of NRT2.1 and the pH_*e*_-dependent phenotype of *nrt2.1* mutant suggest auxiliary functions of the transporter in signaling and regulation of cell elongation. Our data further suggest that pH_*e*_ governs substrate specificity of ABC-type transporters, possibly negotiating growth-defense tradeoffs. Adaptations to the external hydrogen concentration do not appear to be restricted to cells and tissues directly exposed to the soil solution. Instead, the information on pH_*e*_ perceived by root cells appears to systemically propagate and to trigger surprisingly complex changes in the proteome and phosphoproteome of leaf cells. An exciting finding is the putative involvement of TPC in the acclimation to changes in pH_*e*_, possibly regulating the (pH_*e*_- dependent) turnover of cargo proteins involved in signaling or regulating pH_*cyt*_ and pH_*apo*_.

Surprisingly, only very subtle (albeit robust) changes were observed regarding the uptake and homeostasis of mineral nutrients, the availability of which is strongly affected by soil pH (i.e., iron and zinc), suggesting that nutrient and pH signaling run largely separate courses. While our proteomic survey represents just an initial stage in setting the stage to address the question as to how information on pH_*e*_ is transduced into adaptive responses, our data map previously uncharted territory that awaits further experimental exploration.

## Abbreviations —

AA: Acetic Acid
ACN: Acetonitrile
ABC: ATP-Binding Cassette
AHA: H(+)-ATPase
AMT: Ammonium Transporter
ANOVA: Analysis of Variance
CAA: 2-Chloroacetamide
CPKs: Calcium-Dependent Protein Kinases
CalB: Calcium-dependent lipid or phospholipid binding
COPT5: Copper Transporter 5
DEPs: Differentially Expressed Proteins
DPPs: Differentially Expressed Phosphoproteins
DGR2: Duf642 L-Gall Responsive Gene 2
EXL2: Exordium Like 2;
FeEDDHA: Ethylenediamine di-2-hydroxyphenyl acetate ferric
IRT1: Iron-Regulated Transporter 1
IBA: Indole-3-butyric acid
IMAC: Immobilized Metal Affinity Chromatography
KEGG: Kyoto Encyclopedia of Genes and Genomes
LRR: Leucine-Rich Repeat
LSD: Least Significant Difference
NRT: Nitrate Transporter
PAMP: Pathogen-Associated Molecular Patterns
PDR transporter: Pleiotropic Drug Resistance
PGIP1: Polygalacturonase-Inhibiting Protein
PILS5: PIN-Like 5
pFe: free Fe concentration
pH_*apo*_: apoplastic pH
pH_*cyt*_: cytosolic pH
pH_*e*_: environmental pH
RALF: Rapid Alkalization Factor
RGF: Root meristem Growth Factor
SD: Standard Deviation
SEM: Standard Error Mean
SERK: Somatic Embryogenesis Receptor Kinase
SLAH3: SLAC1 HOMOLOGUE 3
TASHTPLATE: Associated SH Domain Containing Protein
TCEP: Tris 2-carboxyethyl phosphine hydrochloride
TEAB: Triethylammonium bicarbonate
TFA: Trifluoroacetic acid
TMT: Tandem Mass Tag
TMK1: Transmembrane Kinase 1
TPC: TPLATE Complex;
VIT: Vacuolar Iron Transporter;
VTL: Vacuolar Iron Transporter Like
ZIP3: Zinc Transporter 3

## Acknowledgments

We sincerely thank Yi-Fang Tsay for providing *nrt2.1* seeds, Chuan-Chih Hsu from the IPMB Proteomics Core Laboratory for their help with proteomic data analysis and Wendar Lin from the Bioinformatic Core Laboratory for help with data mining. Proteomic mass spectrometry analyses, in-gel digestion, and label-free quantification were performed by the Proteomics Core Laboratory sponsored by the Institute of Plant and Microbial Biology and the Agricultural Biotechnology Research Center, Academia Sinica. We acknowledge the use of the BioRender software (https://biorender.com) for generating the figures of this article.

## DATA AVAILABILITY

The mass spectrometry proteomics data have been deposited to the ProteomeXchange Consortium via the PRIDE (Perez-Riverol et al., 2022) (85) partner repository with the dataset identifier PXD039497.

Username

Password: E7PxIs3O

## Supplemental data

This article contains supplemental data.

## Funding and additional information

This work was funded by an Academia Sinica Investigator Award (AS-IA-111-L03) to W.S.

## Author contributions

W.S. original draft and funding acquisition; D.J. experiments and figures. W.S., D.J. data analysis and manuscript editing.

## Supplementary Figures

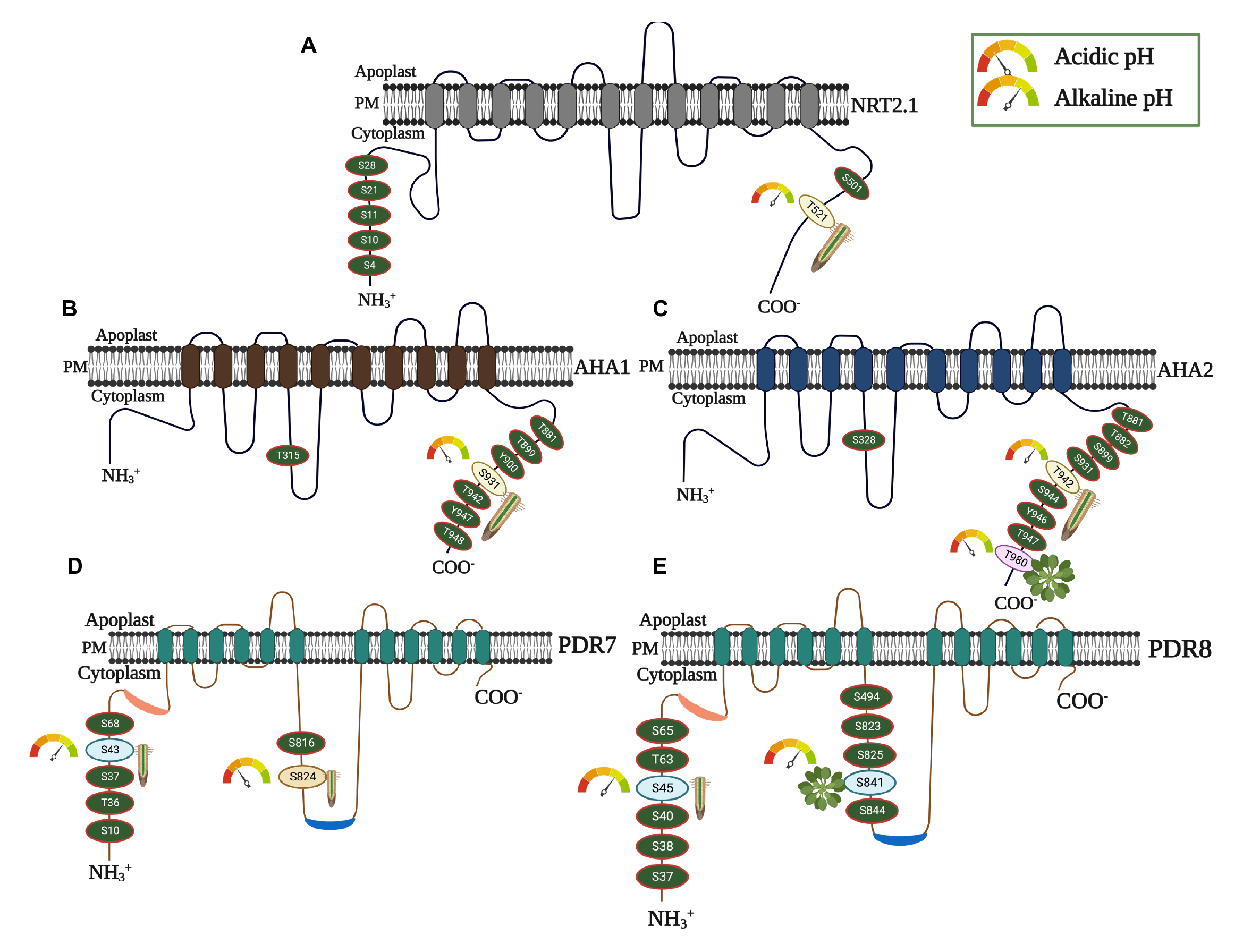

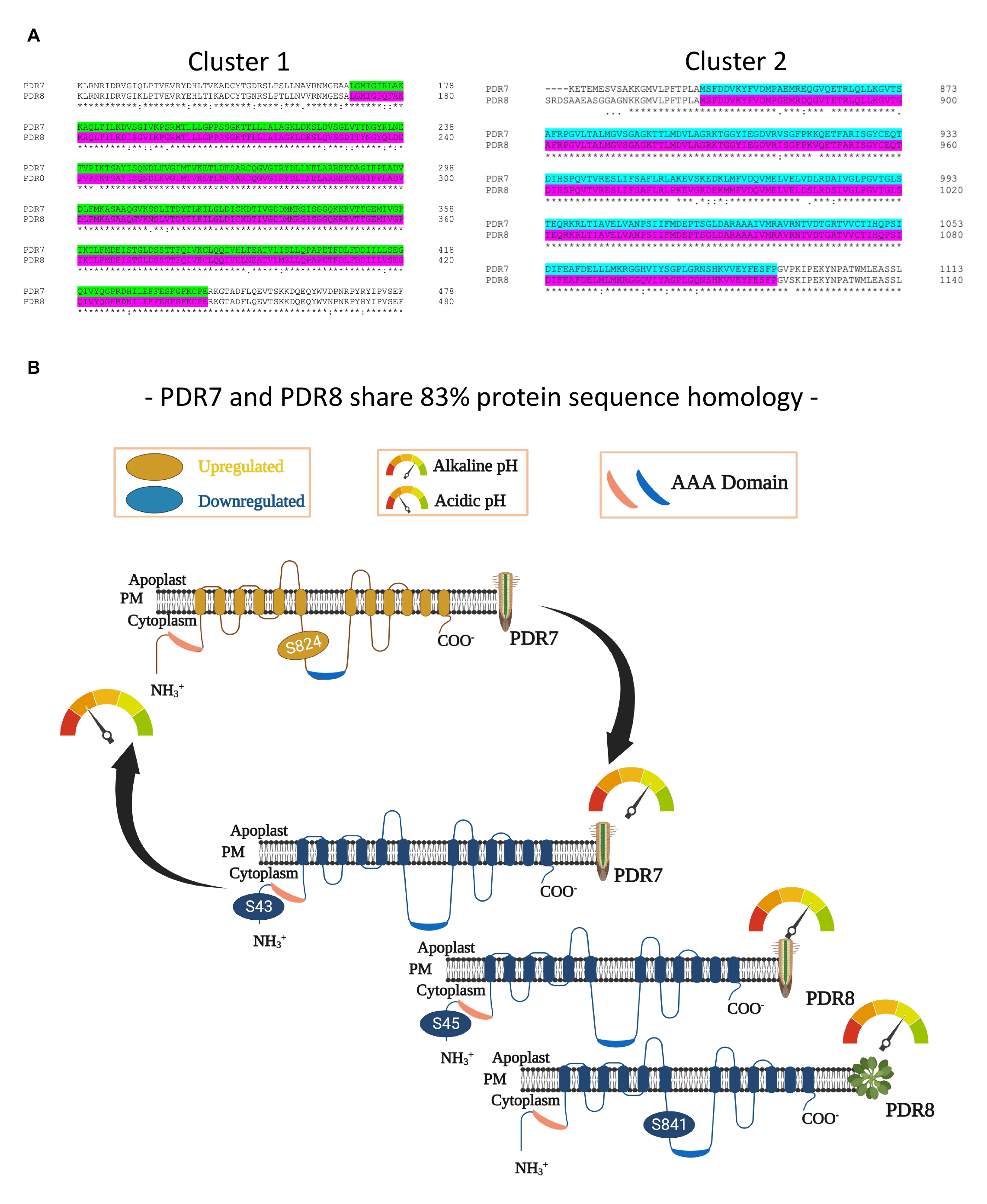

